# Aggregation of *recount3* RNA-seq data improves inference of consensus and tissue-specific gene co-expression networks

**DOI:** 10.1101/2024.01.20.576447

**Authors:** Prashanthi Ravichandran, Princy Parsana, Rebecca Keener, Kaspar D. Hansen, Alexis Battle

## Abstract

**Background:** Gene co-expression networks (GCNs) describe relationships among expressed genes key to maintaining cellular identity and homeostasis. However, the small sample size of typical RNA-seq experiments which is several orders of magnitude fewer than the number of genes is too low to infer GCNs reliably. *recount3*, a publicly available dataset comprised of 316,443 uniformly processed human RNA-seq samples, provides an opportunity to improve power for accurate network reconstruction and obtain biological insight from the resulting networks.

**Results:** We compared alternate aggregation strategies to identify an optimal workflow for GCN inference by data aggregation and inferred three consensus networks: a universal network, a non-cancer network, and a cancer network in addition to 27 tissue context-specific networks. Central network genes from our consensus networks were enriched for evolutionarily constrained genes and ubiquitous biological pathways, whereas central context-specific network genes included tissue-specific transcription factors and factorization based on the hubs led to clustering of related tissue contexts. We discovered that annotations corresponding to context-specific networks inferred from aggregated data were enriched for trait heritability beyond known functional genomic annotations and were significantly more enriched when we aggregated over a larger number of samples.

**Conclusion:** This study outlines best practices for network GCN inference and evaluation by data aggregation. We recommend estimating and regressing confounders in each data set before aggregation and prioritizing large sample size studies for GCN reconstruction. Increased statistical power in inferring context-specific networks enabled the derivation of variant annotations that were enriched for concordant trait heritability independent of functional genomic annotations that are context-agnostic. While we observed strictly increasing held-out log-likelihood with data aggregation, we noted diminishing marginal improvements. Future directions aimed at alternate methods for estimating confounders and integrating orthogonal information from modalities such as Hi-C and ChIP-seq can further improve GCN inference.

## Background

Critical cellular processes including the maintenance of cellular identity, homeostasis, and the cellular response to external stimuli are orchestrated through complex transcriptional co-regulation of multiple genes [1–4]. Gene Co-expression Networks (GCNs) are a commonly used framework to describe gene-gene relationships and are comprised of nodes that represent genes and edges linking co-expressed genes [5]. A comprehensive catalogue of gene co-expression relationships has the potential to characterize genes with unknown functions [6], identify regulatory genes [7], determine changes in regulatory mechanisms that are key to cellular identity [8], and prioritize genes that drive phenotypic variability [9]. Despite the utility of gene networks in understanding biological systems, network inference is still a challenging problem and suffers from both false positive and negative edges [10]. In particular, the typical sample size of most RNA-seq studies is orders of magnitude smaller than the number of gene pairs over which regulatory relationships are inferred, making network inference an underdetermined problem. Additionally, factors such as the stochastic nature of gene expression, experimental noise, missing data, and unobserved technical confounders make it difficult to avoid false positives or negatives. Since the number of possible gene-gene interactions scales with the square of the number of genes examined, a potential solution to increase statistical power by reducing network complexity has been to infer modules or groups of co-expressed genes that are regulated by one or more transcription factors rather than individual gene interactions [11]. While this approach has been successful at decreasing the number of hypotheses tested, and thereby increasing statistical power [12], it does not identify detailed network structure or distinguish between direct and indirect gene interactions. In contrast, network inference by graphical lasso [13, 14] results in the identification of pairwise edges reflecting direct effects, such that the absence of an edge implies the conditional independence of the genes when all other genes are observed. Further, the formulation of graphical lasso enables flexible penalization based on network density which aids in the identification of a network structure that improves the discovery of true gene-gene interactions while reducing false positives [15]. Here, we focus on improving the statistical power of network inference by significantly increasing the number of samples used in network inference, leveraging large-scale publicly available and uniformly processed RNA-seq data from *recount3* [16] which includes human RNA-seq samples from GTEx [17], TCGA [18], and SRA [19, 20].

Since the *recount3* project consists of RNA-seq data compiled from diverse data sources and multiple tissues with inconsistent sample characteristics, we developed a data pre-processing pipeline to identify and exclude outliers, harmonize the observed gene expression, and cluster samples into meaningful biological contexts such that we can reliably infer GCNs. We utilized 95,280 samples following quality control and pre-processing to infer three consensus networks: a universal network, a cancer network, and a non-cancer network, in addition to 27 context-specific networks which included 7 novel contexts only found in *recount3*. We compared strategies for confounder adjustment and data aggregation and found that accounting for confounders in each study followed by the weighted aggregation of covariance matrices by prioritizing studies with more samples resulted in networks with better generalizability and increased ability to recapitulate known biological gene-gene relationships. We observed that context-specific networks constructed using a combination of SRA and GTEx samples outperformed tissue-specific networks inferred solely using GTEx samples at recapitulating known tissue-specific protein-protein interactions (PPIs) and assigning tissue-specific transcription factors to central nodes, demonstrating the value of data aggregation in GCN inference. Furthermore, using S-LDSC [21] we observed that our context-specific networks had significant heritability enrichment attributed to network features when examining traits related to the tissue context. In conclusion, our work provides a carefully annotated RNA-seq data set, outlines best practices for GCN inference by leveraging publicly available RNA-seq data, and a set of consensus and context-specific networks that will aid the scientific community in achieving the full potential of GCN inference in biomedical research.

## Results

### A. Manual annotation and clustering of RNA-seq data from *recount3* identified 27 unique biological contexts

We downloaded uniformly processed RNA-seq samples from humans using the *recount3* R package [16] comprised of experiments from three data sources, The Sequence Read Archive (SRA) [19, 20], Genotype-tissue Expression (GTEx version 8) [17], and The Cancer Genome Atlas (TCGA) [18] and selected 1,747 projects that included 30 or more samples each. Following quality control **(Methods)**, 95,280 human bulk-RNA sequencing samples remained from 50 GTEx tissues (18,828 samples), 33 TCGA cancer types (11,091 samples), and 884 SRA studies (65,361 samples) **(Fig. 1 A)**. The aggregated data includes samples from a wide array of tissues, cell types, and diseases. While GTEx and TCGA studies included metadata specifying the tissue of origin and disease status for all samples, SRA studies had inconsistent nomenclature. Therefore, to obtain reliable labels for SRA samples, we manually parsed sample descriptions to obtain sample characteristics corresponding to tissue type and disease status for 65,361 SRA samples **(Methods)**. Based on curated annotations, 93.5% of TCGA samples and 30.4% of SRA samples were cancer. In contrast, all GTEx samples were non-cancer, as expected **(Fig. 1 B)**. Tissue labels with the greatest number of samples across all three data sources included blood, central nervous system, breast, skin, and lung **(Fig. 1 C)**. SRA included 224 distinct tissue labels derived from manual annotation that was not observed in GTEx or TCGA, and reflected a wide range of disease states including Type I Diabetes, Alzheimer’s disease, bipolar disorder, arthritis, cancer, and infectious conditions **(Fig. 1 D)**. We grouped SRA samples based on their study accession IDs, GTEx samples by tissue, and TCGA samples by cancer code (**Methods**). To simplify terminology, we defined each group of samples from a data source as a single study. To leverage the extensive biological diversity in the data, we inferred two broad types of networks: consensus and tissue context-specific (context-specific). Our universal consensus network included all samples, regardless of tissue or disease. Our non-cancer consensus network included healthy samples and samples with disease status other than cancer. Finally, our cancer consensus network solely included cancerous samples. We restricted our context-specific networks to non-cancerous samples grouped by tissue context. We did not examine differential coregulation resulting from non-cancer disease, and regressed these effects from gene expression. Thus, by including SRA, *recount3* provides an unprecedented opportunity to examine unique contexts that were not previously studied in GTEx and TCGA.

**Fig. 1.**
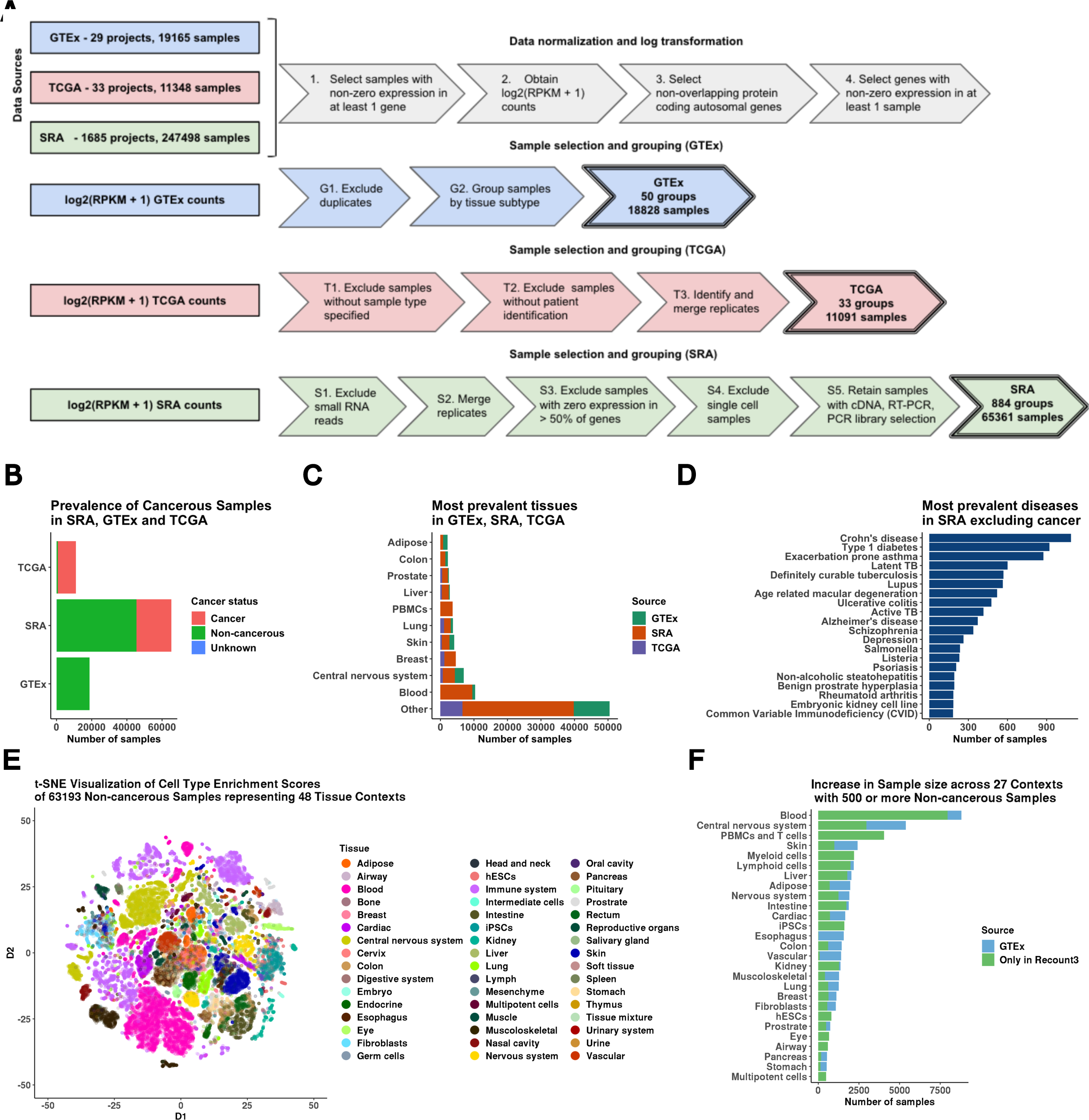
Overview of data pre-processing and annotations. **(A)** Gene expression data was RPKM normalized and log-transformed along with gene-specific and sample-specific filters. Based on the data source, normalized gene expression was processed to merge replicates, and exclude miRNA and scRNA seq samples. **(B)** Number of samples which were annotated to be non-cancer, cancer, and unknown based on available metadata across GTEx, SRA, and TCGA. **(C)**Top 10 tissue labels by sample size across all three data sources: SRA, GTEx, and TCGA. **(D)**Top 20 diseases by sample size found in SRA that are not cancer. **(E)** t-SNE projection of xCell deconvolution scores of 63,193 non-cancerous samples colored by the tissue of origin. **(F)** Increase in the sample size of 27 tissue contexts by using SRA samples compared to GTEx only. SRA studies included 7 novel contexts which were not available in GTEx.

Across the three data sources, SRA, GTEx, and TCGA, we obtained 266 unique manually annotated tissue labels with a median sample size of 31 which was much lower than the number of protein-coding genes. Therefore, we used a study-pooling strategy based on related tissue context to increase power **(Methods)**. Mapping manual annotations to tissue contexts **(Additional File 1: Supp. Table I)** generated 48 tissue contexts across 63,193 non-cancerous samples for context-specific network analysis. In each context, to ensure that a tissue context represented samples with similar cell-type composition as estimated by xCell [22] deconvolution, we learned a joint lower-dimensional t-SNE embedding using cell-type deconvolution scores, and for 25 contexts with more than 500 samples before outlier exclusion, we detected and excluded outliers **(Additional File 2: Supp. Fig 1)**. For the immune context, we observed that samples displayed extensive heterogeneity in cell-type composition. Thus we further separated this group into B cells, PBMCs/ T cells, and myeloid cells **(Additional File 2: Supp. Fig 2)**.

In total, we obtained 27 contexts including 7 tissue contexts that were not present in GTEx including airway, eye, human embryonic stem cells, induced pluripotent stem cells, multipotent cells, myeloid cells, and PBMCs / T cells **(Fig. 1 E)**. The number of samples present in each tissue context following outlier exclusion varied from 8,797 (Blood) to 485 (Multipotent cells) **(Fig. 1 F)**. Further, for tissues that were present in GTEx, the ratio of the number of samples across all data sets to those in GTEx only ranged from 14.13 (Kidney) to 1.01 (Esophagus) **(Fig. 1 F)**. These harmonized samples have increased context resolution, i.e. include novel contexts which were not examined in GTEx, and increased sample size which can be used to improve the inference of consensus and context-specific networks.

### B. Data aggregation improves the inference of consensus and context-specific GCNs

The median study-specific sample size across the three data sources was 44 for SRA, 309 for TCGA, and 285 for GTEx. Further, 766 of 884 SRA studies had a sample size lesser than 100. PC-based data correction has been used within a single study to reduce potential false positive gene regulatory associations [23, 24], but best practices for applying PC-based data correction in the context of aggregating multiple studies to infer GCNs have not been fully examined. First, we sought to determine whether data correction should be performed jointly across all samples, across samples belonging to a specific tissue context and multiple studies, or across samples from a specific tissue context and a specific study. We observed that PCs recapitulate different sample characteristics depending on the level at which data aggregation is performed. PCs calculated across all samples were driven predominantly by tissue labels followed by technical confounders (e.g. study and data source) **(Additional File 2: Supp. Fig 4 A-D, Supp. Fig 5 A)**. PCs which were calculated across samples belonging to a single tissue context (blood) but multiple studies were predominantly driven by study and data source. Further, accounting for tissue heterogeneity enabled us to better model cancer status and disease annotations using PCs **(Additional File 2: Supp. Fig 4 E-H, Supp. Fig 5 B)**. Finally, when we limited samples to a specific tissue context and a specific study (GTEx-skeletal muscle), we found that the top PCs were significantly associated with technical batch, consistent with the findings of Parsana et al. [24] **(Additional File 2: Supp. Fig 4 I-L, Supp. Fig 5 C)**. Thus regressing PCs computed by accounting for tissue and cross-study heterogeneity from expression is integral to excluding technical effects and unwanted biological signals.

In addition to comparing the effect of PC-based data correction, before and post-aggregation, where the number of PCs is selected based on the permutation method described by Buja et. al [25] and Leek et. al [23], we sought to examine the consequences of aggregating empirically estimated covariance matrices considering all studies equally (unweighted aggregation) as compared to weighting covariance matrices from studies with a greater number of samples more (weighted aggregation) on GCN inference. Thus, we used 4 paradigms: (1) Aggregating data before PC-based data correction followed by estimation of empirical covariance from residual expression, (2) PC-based data correction applied to individual studies followed by aggregation of residual expression and joint estimation of empirical covariance, (3) Unweighted aggregation of covariance matrices inferred from each study separately after study-specific PC-based correction and (4) Weighted aggregation of covariance matrices computed from individual studies following study-specific PC-based data correction, where the weights were the ratio of the study sample size to the total number of samples used in network reconstruction **(Fig. 2 A, Methods)**.

**Fig. 2.**
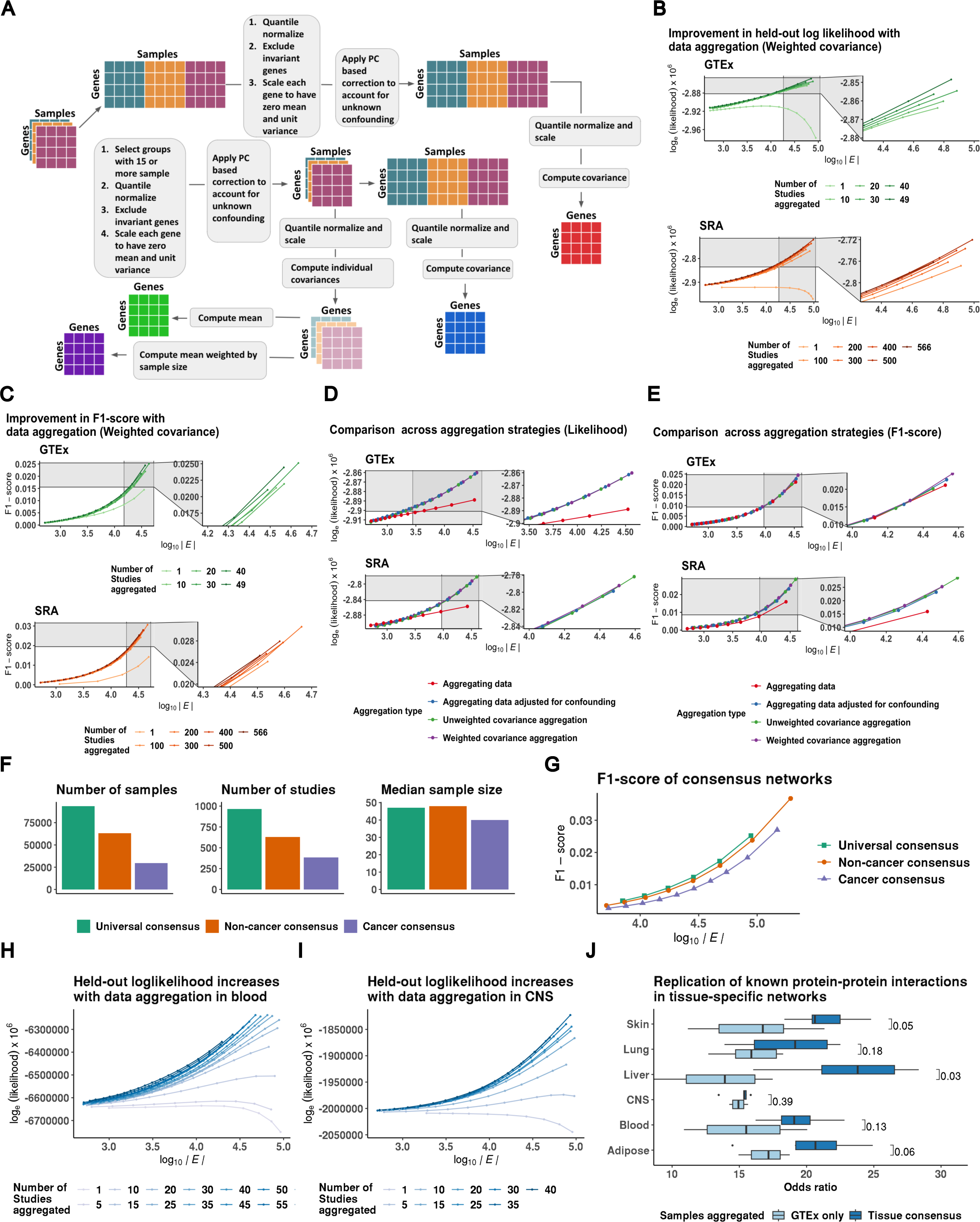
Comparison of aggregation strategies to optimize network reconstruction. **(A)** Outline of strategies to compare data correction before and after aggregation and weighted and unweighted aggregation of single tissue/ study covariance matrices included (1) Aggregating data before PC-based data correction followed by estimation of empirical covariance from residual expression (Aggregating data, orange), (2) PC-based data correction applied to individual studies followed by aggregation of residual expression and joint estimation of empirical covariance (Aggregating data adjusted for confounding, brick red), (3) Unweighted aggregation of covariance matrices inferred from each study separately after study-specific PC-based correction (Unweighted covariance aggregation, purple) and (4) Weighted aggregation of covariance matrices computed from individual studies following study-specific PC-based data correction (Weighted covariance aggregation, magenta). **(B)** Held-out log-likelihood of networks inferred by sequentially aggregating either 10 GTEx studies or 100 SRA studies at a time. **(C)** F1-score of networks inferred by sequentially aggregating either 10 GTEx studies or 100 SRA studies at a time when compared to canonical pathways compiled from KEGG, Biocarta, and Pathway Interaction Database. **(D)** Comparison of held-out log-likelihood corresponding to networks inferred over 49 GTEx studies or 566 SRA studies using four different aggregation strategies including aggregating data, aggregating data adjusted for confounding, unweighted, and weighted aggregation of covariance matrices. **(E)** Comparison of F1-scores of obtaining edges corresponding to canonical pathways from KEGG, Biocarta, and Pathway Interaction Database in networks inferred over 49 GTEx studies or 566 SRA studies using four different aggregation strategies including aggregating data, aggregating data adjusted for confounding, unweighted, and weighted aggregation of covariance matrices. **(F)** Total number of samples, number of individual studies, and the median sample size of each study which were used in the inference of universal consensus, non-cancer consensus, and cancer consensus networks. **(G)** Comparison of F1-scores of obtaining edges corresponding to canonical pathways in the three consensus networks, universal, non-cancer, and cancer, across networks with density (*E*) varying between 5e3 to 5e6 edges. **(H)** Log-likelihood of GTEx blood samples based on networks inferred by sequentially aggregating SRA blood studies five at a time for densities ranging from 1e3 to 1e5 edges. **(I)** Log-likelihood of GTEx CNS samples based on networks inferred by sequentially aggregating SRA CNS studies five at a time for densities ranging from 1e3 to 1e5 edges. **(J)** Odds ratio of finding edges corresponding to tissue-specific protein-protein interactions (PPIs) derived from SNAP in tissue-context-specific networks inferred using all available samples vs. only samples found in GTEx for six tissue contexts.

To compare strategies, we split non-cancerous samples into two data splits, GTEx and SRA **(Additional File 2: Supp. Fig 6)**, followed by network inference by graphical lasso on one of the two data splits and evaluation with the held-out split by computing the held-out log-likelihood. Details pertaining to the number of studies, samples and median PCs regressed for incremental data aggregation are provided in **Additional File 3: Supp. Table II**. Additionally, we assessed the recapitulation of known biological pathways by computing the F1-score of finding canonical gene-gene relationships compiled from KEGG [26], Biocarta, and Pathway Interaction Database [27] obtained using Enrichr [28, 29] **(Additional File 4: Supp. Table III)**. Paradigms in which PC-based data correction preceded aggregation led to a strict increase in held-out log-likelihood and F1-scores of known gene relationships from canonical pathways with the addition of more studies **(Fig. 2 B, C, Additional File 2: Supp. Fig 7, 8)**. This suggests that data aggregation resulted in GCNs with greater generalizability and recapitulated known biology better when technical confounders were estimated and regressed for individual studies. Since data aggregation led to a decrease in the network density for a particular value of the penalization parameter **(Additional File 2: Supp. Fig 9)**, we tested whether estimating denser networks would result in higher held-out log-likelihood specifically when the networks were inferred over a greater number of samples, but instead observed that higher densities led to overfitting and lower generalizability **(Additional File 2: Supp. Fig 10)**. Further, the marginal improvement in network reconstruction diminished with the subsequent rounds of aggregation. While all methods that estimated technical confounders within each study performed similarly and were superior to estimating technical confounders across all samples when evaluated by held-out log-likelihood **(Fig 2 D)**, we found that weighted aggregation of covariance matrices led to a slight improvement in the F1-score of the networks when compared to canonical pathways **(Fig. 2 E)**, suggesting that this is the optimal strategy among the alternatives compared.

We inferred consensus GCNs across diverse tissues by weighted aggregation of covariance matrices estimated from residual expression and graphical lasso **(Methods)**. In addition to a universal consensus network which was inferred across 966 studies with sample size greater than or equal to 15, amounting to 95,276 samples across 48 tissue contexts, we constructed a non-cancer consensus network and a cancer consensus network. The non-cancer consensus network reflects data aggregated across 629 studies and 63,031 samples, and the cancer network reflects 386 studies and 29,967 samples **(Fig. 2 F)**. We evaluated each consensus network to recapitulate previously reported gene-gene interaction using the F1 score. Across a range of network densities, we obtained a higher estimate of the F1 score from the universal consensus network, followed by non-cancer and cancer consensus networks, which mirrors differences in sample size **(Fig. 2 G)**.

We inferred networks across 27 tissue contexts and examined the impact of data aggregation on context-specific network reconstruction by considering GTEx samples (GTEx only) or by aggregating across samples from all data sources for that tissue context. The number of samples, studies, and median study-specific sample size for each tissue context in either aggregation setting are provided in **Additional File 2: Supp. Fig 3**. As in the consensus network inference, we used weighted aggregation of covariance matrices as the input to network inference by graphical lasso **(Methods**). To quantify the improvement in network reconstruction with data aggregation, we examined two tissue contexts with the largest sample size: blood and central nervous system (CNS). In both cases, we sequentially aggregated the blood or CNS SRA studies and computed the held-out log-likelihood utilizing context-matched GTEx samples. Similar to the results obtained from our consensus network analyses, we found that data aggregation led to a strict increase in held-out log-likelihood with additional studies and the greatest increase was observed while aggregating the first 20 studies (blood) and 15 studies (CNS) **(Methods, Fig. 2 H, I)**.

Orthogonally, we verified that our networks recapitulated known context-specific gene relationships by examining the enrichment of previously known tissue-specific protein-protein interactions (PPIs) from SNAP [30] in edges belonging to six tissue context-specific networks (adipose, blood, CNS, liver, lung, and skin) **(Methods**). Data aggregation and increased sample size in the network significantly increased the estimated odds ratio of tissue-specific PPIs in the liver (Median GTEx OR = 7.85, Median Aggregate OR = 38.85, p = 0.03) and skin (Median GTEx OR = 21.83, Median Aggregated OR = 27.09, p = 0.05), and there was weak enrichment in adipose (Median GTEx OR = 18.49, Median Aggregated OR = 21.14, p = 0.06). In other tissue contexts, we observed a higher median odds ratio of PPI enrichment in networks inferred by data aggregation compared to GTEx-only networks **(Fig. 2 J)**. Thus, we demonstrated that data aggregation led to the improved inference of consensus networks that capture ubiquitous biological pathways and tissue context-specific networks that capture tissue biology by observing better network generalizability and reproduction of known biological processes.

### C. Central network nodes are evolutionarily constrained and include genes that are critical to tissue identity

The biological information captured by a GCN can be evaluated by comparing individual network edges, by examining whether there are edges between genes that are known to interact in a particular cellular pathway [31], or by examining the properties of hubs [32–34], network nodes with a high number of connections. Since eukaryotic transcriptional networks typically consist of a subset of genes, often transcription factors, that regulate many downstream target genes [35], we chose specific network densities such that the selected networks are approximately scale-free **(Methods, Additional File 2: Supp. Fig 11; Additional File 5: Supp. Table IV, Additional File 6: Supp. Table V)**. We computed different measures of centrality corresponding to each network node and tested for the enrichment of genes involved in GO terms that reflect ubiquitous or context-specific processes **(Additional File 7: Supp. Table VI)** among network nodes selected with progressively increasing thresholds for degree centrality against a background of all 18,882 protein-coding genes **(Methods)**. Central nodes from all three consensus networks were strongly enriched for genes involved in functions such as microtubule-based process, chromosome organization, and regulation of organelle organization **(Fig. 3 A, Additional File 2: Supp. Fig 12)**. In contrast, we found that central nodes from blood context-specific networks were enriched for genes associated with platelet activation, leukocyte differentiation, and leukocyte chemotaxis to a greater extent than central genes derived from either consensus or a discordant context-specific network (CNS) **(Fig. 3 B)**. Further, these trends were reflected across multiple tissue contexts; context-specific networks corresponding to CNS **(Fig. 3 C)**, skin, liver, and lung were enriched for genes associated with tissue-matched GO terms **(Additional File 2: Supp. Fig 13)**. Thus, while central genes from consensus networks included genes involved in essential cellular processes, context-specific gene relationships are lost with global aggregation and are unlikely to be recovered by increasing the sample size.

Next, we evaluated whether hub genes were evolutionarily constrained and their role in complex traits or diseases **(Methods)**. We first binned network nodes such that genes with no neighbors were assigned to Quintile 1, and those with at least 1 connecting edge were grouped based on estimated quantiles of closeness centrality. Consensus network hub genes (Quintile 5) had a significantly higher excess overlap (>1) with evolutionarily constrained gene sets than peripheral nodes (Quintiles 1-3), using metrics including high pLI genes, high s_het_ genes, and high missense Z-score genes. These trends were similar in context-specific networks with some variation in the strength of the trend across tissues likely driven by sample size differences and differential power in inferring these networks **(Fig. 3 D, Additional File 2: Supp. Fig 15)**. We then examined the enrichment of eQTL-deficient, ClinVar, OMIM, and FDA drug-targeted genes across quintiles of network centrality **(Additional File 2: Supp. Fig 14, 15, Additional File 8: Supp. Table VII)**. We observed that peripheral genes (Quintile 1) and central genes (Quintile 5) of consensus networks exhibited an excess overlap > 1 of eQTL deficient genes (those which lacked significant *cis*-regulatory variants), while genes with intermediate connectivity were depleted of eQTL deficient genes, while the six context-specific networks had widely variable trends. While central genes from consensus networks were weakly depleted for OMIM and FDA drug-targeted genes, we observed an excess overlap > 1 of hub genes belonging to context-specific networks. This could be explained by our earlier observations that central nodes from consensus networks were involved in essential and non-specific cellular processes while central nodes from context-specific networks were both specific and critical to tissue identity, thus, altering central consensus network hub nodes may have widespread and potentially deleterious off-target effects while context-specific hub nodes may identify targetable genes with tissue-specific effects. Finally, we observed that matched tissue-specific transcription factors (TFs) from Pierson et. al [36] **(Additional File 9: Supp. Table VIII)** had a significantly greater number of neighbors than 88 general TFs for context-specific networks corresponding to blood (p < 1e-4), lung (p < 1e-4), and skin (p < 0.01), while in other tissues, except for CNS, the median degree of tissue-specific TFs was greater than general TFs and non-transcription factors **(Methods)**. In contrast, while tissue-specific and general TFs had more neighbors in consensus networks when compared to non-transcription factors, we found no significant differences in the degree distribution of tissue-specific TFs to general TFs in either the universal or non-cancer consensus network **(Fig. 3 E)**. Therefore, while both consensus and context-specific central genes were enriched for genes under high-selection pressure, context-specific hub nodes were more likely to be OMIM genes, drug targets and tissue-specific TFs.

**Fig. 3.**
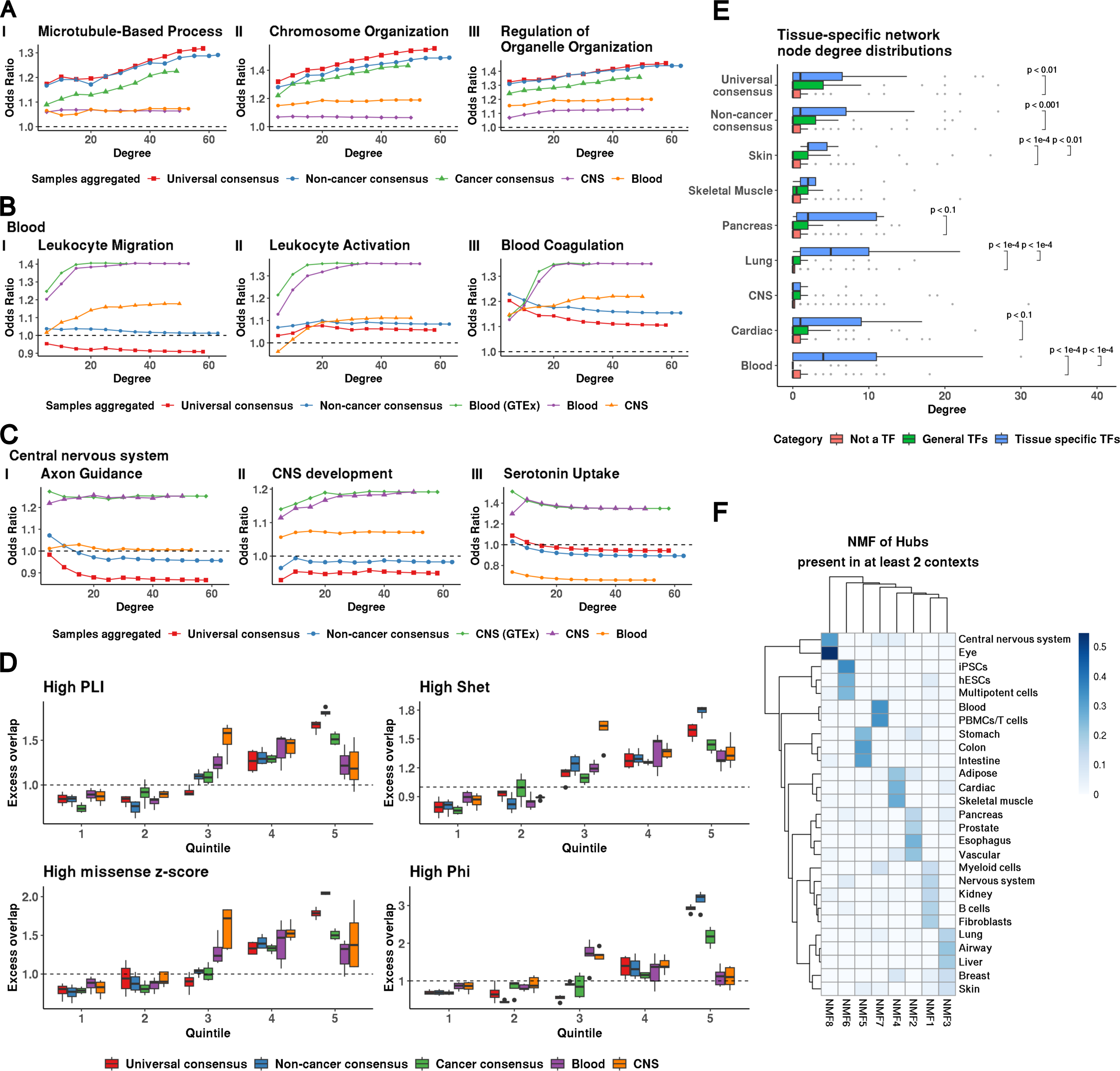
Properties of central network nodes of consensus and context-specific networks. **(A-C)** Enrichment of genes involved in GO processes among network genes selected with increasing thresholds of degree connectivity in three consensus networks, universal (λ = 0.18, 7087 edges), non-cancer (λ = 0.18, 7355 edges), and cancer (λ = 0.24, 7552 edges), as well as two context-specific networks blood (λ = 0.24, 7283 edges) and CNS (λ = 0.28, 8430 edges). Tissue-context-specific networks were inferred only using non-cancerous samples. Blood (GTEx) or CNS (GTEx) networks were inferred using samples only found in GTEx while Blood and CNS networks were inferred using samples from GTEx and SRA. **(D)** Distribution of the excess overlap of evolutionarily conserved gene sets (Methods) for network nodes binned by the number of neighbors (degree) corresponding to universal consensus networks (λ = 0.14, 0.16, 0.18, 0.20), non-cancer consensus network (λ = 0.14, 0.16, 0.18, 0.20), cancer consensus networks (λ = 0.20, 0.22, 0.24, 0.26), blood network (λ = 0.18, 0.20, 0.22, 0.24, 0.26), and CNS network (λ = 0.24, 0.26, 0.28, 0.30, 0.32). Quintile 1 reflects nodes with no neighbors. Nodes with non-zero neighbors are split based on the degree quartile they belong to (Quintiles 2-5). We evaluated the excess overlap of 3,104 loss-of-function (LoF) genes with pLI > 0.9,2,853 genes with a S_het_ > 0.1, 588 genes with a Phi-score > 0.95, and 1,440 genes strongly depleted for missense mutations (high missense z-score). **(E)** The degree distribution of network nodes that are tissue-specific transcription factors (TFs) in blood (52 TFs), lung (58 TFs), skin (10 TFs), pancreas (16 TFs), cardiac (17 TFs), muscle (7 TFs), CNS (51 TFs), general transcription factors (88 TFs), and protein-coding genes which are not transcription factors in universal consensus (λ = 0.18, 7,087 edges), non-cancer consensus (λ = 0.18, 7,355 edges), skin (λ = 0.26, 7,567 edges), skeletal muscle (λ = 0.26, 6,254 edges), pancreas (λ = 0.32, 7,615 edges), lung (λ = 0.30, 6,349 edges), CNS (λ = 0.30, 6,316 edges), cardiac (λ = 0.30, 6,481 edges), blood (λ = 0.24, 7,283 edges). Pairs with no significance reported were not statistically distinct (p > 0.1). **(F)** Factor weights were obtained by non-negative matrix factorization of the presence of hub genes in tissue-specific networks with ∼ 7,000 edges. Details of the penalization parameter λ and density of selected networks for each tissue context are provided in **Supp. Table X**.

We examined similarities and differences between network architectures of the non-cancer consensus and cancer consensus networks based on shared and distinct hub genes, since central genes are likely to be more relevant to network functionality [32], and the identification of hubs has led to the discovery of genes involved in cancer [33, 34], tissue regeneration [37], and other diseases [38, 39]. Specifically, we found 296 shared hubs between cancer (λ = 0.24, 7552 edges) and non-cancer (λ = 0.18, 7,355 edges) consensus networks, and 312 hubs which were specific to the cancer consensus network **(Additional File 10: Supp. Table IX)**. Cancer-specific hub genes were enriched for pathways such as ncRNA metabolic processing (GO:0034660, p = 5.2e-02) which is believed to play a role in metabolic reprogramming in cancer, DNA damage response (GO:0006974, p = 1.17e-12) which has been posited to play a role in cancer cell survival in non-optimal conditions, and response to ionizing radiation (GO:0010212, p = 6.1e-04). Further, we found that cancer-specific hub genes were enriched for a plethora of DNA repair and replication pathways including, double-strand break repair via homologous recombination (GO:0000724, p = 6.36e-06), recombinational repair (GO:0000725, p = 1.35e-06), interstrand cross-link repair (GO:0036297, p = 5.48e-03), and double-strand break repair via break-induced replication (GO:0000727, p = 1.36e-03) **(Additional File 11: Supplementary File 1)**.

To examine whether network properties were shared between related tissues we compared the overlap of hub, we considered all hub genes found in at least two tissue contexts (N=1,956). Grouping tissue contexts based on hub genes using non-negative matrix factorization with eight latent factors **(Methods)** led to the grouping of related tissues such as blood and PBMCs/T cells (Factor 7) **(Fig. 3 F, Additional File 12: Supp. Table X)**. As expected, we found that hub genes that led to the grouping of blood and PBMCs/T cells were enriched for defense response (GO:0006952, p = 2.7-24) and cytokine production (GO:0001816, p = 9.9e-07) **(Additional File 13: Supplementary File 2)**. Further, Factor 6 which led to the grouping of hESCs, iPSCs, and multipotent cells comprised of hub genes which were enriched for gastrulation (GO:0007369, p = 5.9e-04) and circulatory system development (GO:0072359, p = 6.2e-03) **(Additional File 14: Supplementary File 3)**.

### D. Genes with high network centrality are proximal to variants enriched for complex trait heritability

Previous work by Kim et al. [40] reported that network topology annotations did not contribute to heritability once the LDSC baseline model [21] was included. We examined whether data aggregation would increase the utility of network features for heritability analysis independently of baseline functional annotations. We meta-analyzed estimates of heritability enrichment, the ratio of the proportion of heritability explained by SNPs belonging to an annotation to the proportion of SNPs in the annotation, and τ^*^, an estimate of the heritability of SNPs unique to the annotation [21], using a random effects model to obtain a summary of effect sizes estimated for a set of 42 independent traits considered by Kim et al. **(Methods, Additional File 15: Supp. Table XI)**. We estimated both heritability enrichment and τ^*^ by either conditioning an annotation corresponding to whether a variant was located in a 100 kilobase window of all protein-coding genes (all-genes annotation), or conditioning on 97 functional annotations such as known enhancer and promoter regions which are included in the baseline-LD model and the all-genes annotation (all-genes + baseline). Similar to the results found by Kim et al., we observed a significant estimate τ^*^ corresponding to our consensus network-derived annotations when conditioning on just the all-genes annotation, however, when conditioning on the baseline-LD model, the τ^*^ observed for consensus network-derived annotations were no longer significant **(Fig. 4 A)**. While we found no significant differences in enrichment across network density when conditioned on the all-genes annotation, we observed that τ^*^ decreased with a decrease in network density. There were no significant differences in either enrichment or τ^*^ with network density when conditioning on the baseline-LD annotations **(Additional File 2: Supp. Fig 16)**. Further, our observations were not dependent on the traits studied and remained consistent when we applied s-LDSC to 219 UKBB traits **(Additional File 16: Supp. Table XII; Additional File 2: Supp. Fig 17)**

**Fig. 4.**
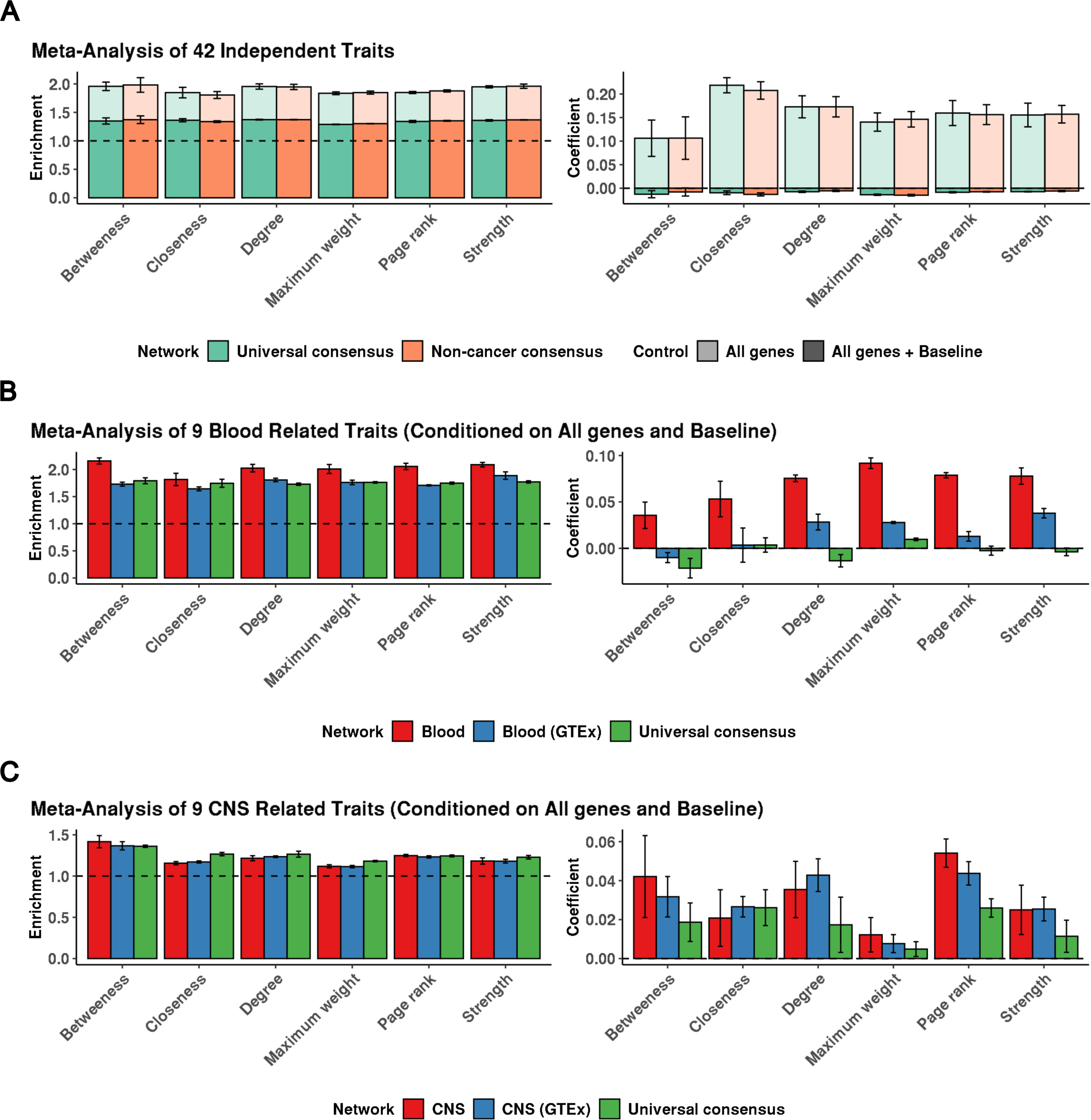
Heritability enrichment of network annotations. Mean and standard deviation of heritability enrichment and the coefficient τ^*^, an estimate of the heritability of SNPs unique to the annotation. All genes: whether a variant was located in a 100 kilobase window of all protein-coding genes, All genes + baseline: all-genes annotation in addition to 97 functional annotations such as known enhancer and promoter regions. **(A)** Meta-analysis of 42 independent traits for six centrality measures obtained from the universal consensus network and non-cancer consensus networks corresponding to values of the penalization parameter *λ* between 0.14 - 0.20. **(B)** Meta-analysis of 9 blood-related traits including, Crohn’s disease, Ulcerative colitis, Rheumatoid arthritis, Allergy Eczema, Eosinophil count, Red blood cell count, White blood cell count, Red blood cell width, and Platelet count for network annotations from Blood GTEx (*λ* = 0.24 - 0.32), Blood consensus (*λ* = 0.18 - 0.26), and Universal consensus network (*λ* = 0.14 - 0.20). **(C)** Meta-analysis of 9 CNS-related traits including, Alzheimer’s disease, Epilepsy, Parkinson’s disease, Bipolar disorder, Smoking cessation, Waist-hip-ratio adjusted BMI, Schizophrenia, Major depressive disorder, and Number of alcoholic drinks per week, for network annotations corresponding to 6 centrality measures derived from CNS GTEx (*λ* = 0.26 - 0.32), CNS (*λ* = 0.20 - 0.28), and Universal consensus network(*λ* = 0.14 - 0.20).

A possible explanation for the lack of heritability enrichment signal unique to network annotations is the redundancy between the baseline LD annotations and network topology annotations. Therefore, we hypothesized that context-specific data aggregation could prioritize variants enriched for heritability of concordant traits independent of baseline annotations. We applied s-LDSC to network centrality annotations derived from networks inferred only from GTEx blood samples (blood GTEx), networks inferred by aggregating *recount3* blood samples (blood), and as a control, networks inferred by aggregating all samples (universal consensus), for a subset of 9 blood-related traits from the 42 independent traits (Crohn’s disease, rheumatoid arthritis, ulcerative colitis, eosinophil count, platelet count, red blood cell count, red blood cell width, white blood cell count, and eczema) **(Additional File 17: Supp. Table XIII)**. As with the consensus networks, tissue-specific networks displayed similar trends in heritability estimates with network density **(Additional File 2: Supp Fig 18, 19)**. When we conditioned on the baseline-LD annotations, we observed that annotations derived from the blood consensus networks had a significant τ^*^ across all centrality annotations, while blood GTEx networks had a significant τ^*^ for strength, degree, maximum weight and page rank **(Fig. 4 B)**. In contrast, we did not observe a significant τ^*^ corresponding to annotations derived from the universal consensus network. We examined the generalizability of our results by conducting a similar experiment in CNS samples, another tissue with a large sample size. We applied s-LDSC to annotations derived from CNS networks inferred from GTEx samples (CNS GTEx), CNS networks inferred by aggregating samples from *recount3* (CNS consensus), and the universal consensus networks for CNS-related traits which included waist-hip ratio adjusted BMI from the earlier set of 42 traits, as well as 7 traits from the Psychiatric Genomics Consortium (Alzheimer’s, epilepsy, Parkinson’s, bipolar disorder, smoking cessation, schizophrenia, major depressive disorder, and number of alcoholic drinks per week) **(Additional File 18: Supp. Table XIV)**. We found that annotations derived from both universal consensus and CNS-specific networks led to significant non-zero τ^*^ when conditioning on the baseline-LD model. While we note that significant non-zero τ^*^ was observed for the consensus networks for the chosen set of CNS traits in contrast to the 42 independent traits, possibly due to power, study quality, and other attributes of the GWAS, we found that annotations from tissue-specific networks led to significantly higher estimates of τ^*^ and outperformed consensus networks **(Fig. 4 C)** for all centrality measures except closeness. Further, for betweenness, maximum weight, and page rank centrality, CNS consensus networks outperformed CNS GTEx networks, similar to the results in blood, demonstrating context-specific data aggregation results in network annotations that are enriched for trait heritability across tissue contexts. Across both sets of blood- and CNS-related traits, we found that page rank centrality-derived annotations, which captured both the number of connections that a node has in addition to the centrality of its neighbors to determine the importance of a connection, performed consistently well. We conclude that context-specific aggregation results in identifying network central genes that are enriched for the heritability of concordant traits, and an increased sample size leads to a greater heritability enrichment signal.

## Discussion

GCNs aid in determining changes in regulatory mechanisms that are key to cellular identity and prioritizing genes that drive phenotypic variability. However, conventional network analyses are often too underpowered to reliably discover gene-gene relationships and are compromised by spurious false positives and false negatives that result from limited power, noise, and unobserved technical confounders. We leveraged publicly available RNA-seq data from *recount3* and manually curated tissue/cell type annotations to improve the inference of consensus and context-specific GCNs. Utilizing data splits, we demonstrated that accounting for confounders within individual studies followed by weighted aggregation of empirical covariance matrices led to the best improvement in network characteristics with data aggregation across multiple paradigms.

We then inferred three consensus networks (universal, non-cancer, and cancer networks) that recapitulated ubiquitous biological processes. Further, we aggregated data belonging to individual tissue contexts to infer 27 tissue context-specific networks that were enriched for matched tissue-specific PPIs and shared similarities across related tissues. All networks and sample annotations are made publicly available as a resource for future studies.

Central genes from both consensus and context-specific networks were enriched for high PLI, and high Phi genes, indicating that hub genes are enriched for genes under high selective pressure. Context-specific hub genes were enriched for FDA-approved drug targets and OMIM genes while central genes from consensus networks which were inferred over a greater number of samples were depleted for both categories. Thus, context-specific information was lost by global aggregation, cannot be recovered by data aggregation or increased sample sizes, and is important to identifying drug targets and disease mechanisms. While network central genes determined by global data aggregation in the consensus network did not explain trait heritability independent of known functional annotations in the baseline-LD model, we found that context-specific data aggregation prioritized variants enriched for concordant trait heritability that did not overlap with previously known functional annotations. Thus topological properties of genes from context-specific GCNs hold significant promise as a functional annotation for identifying genetic variation that contributes to complex trait heritability.

A commonly used approach to identify genes associated with complex traits is to use colocalization analysis between GWAS and eQTL studies, however, often only about half of the signals colocalize with an eQTL [41]. Recent work by Mostafavi et al [41] demonstrated that genes driving GWAS signals were often genes with complex context-dependent regulatory architecture and were depleted for eQTL variants. This has raised a call in the computational genomics community for orthogonal approaches to identify genes involved in complex traits. We found that annotations derived from context-specific GCNs are informative of trait heritability independent of context-agnostic functional annotations. This suggests that tissue- and context-specific network centrality and other network properties could be used to help prioritize genes near GWAS loci [42] or supplement eQTL data.

One of the major challenges in network inference remains the presence of unobserved technical confounders and undesirable biological signals which leads to spurious network edges and precludes causality claims. While PC-based data correction has been extensively utilized to reduce false positives resulting from confounding, recent work by Cote et. al [43] suggests that PC-based data correction, and related methods such as PEER [44] and CONFETI [45], may over-correct expression data and remove biological co-expression of potential interest. Correcting or modeling confounders is essential to network accuracy, so tuning parameters such as the number of latent factors to correct as well as exploring alternative methods will continue to be important. Alternate approaches to handle confounding and infer causal regulatory relationships include instrumental variable analysis through the construction of local genetic instruments as outlined by Lujik et al. [46], however, since central network nodes are evolutionarily constrained and tightly regulated, it can be challenging to construct well-tracking genetic instruments for central genes,. Publicly available RNA-seq data including *recount3*, the extensive annotations we provide, and recent work which illustrated genotype calling using RNA-seq data [47] could improve our ability to detect context-specific cis-regulatory effects, the reconstruction of local genetic instruments, and hence causal regulatory network inference.

Future directions aimed at improving GCN inference could leverage our extensively annotated sample characteristics and data aggregation strategies with complementary strategies including sharing information between related contexts [48] to increase the effective sample size, introducing constraints or priors corresponding to known regulatory relationships [49], and using alternate statistical measures of expression similarity that capture non-linear associations between genes [50]. Additionally, heuristic algorithms such as the one proposed by Opgen-Rhein et al. [51] could be utilized to enrich our current networks with directionality information. Finally, while we studied tissue-contexts, we provide annotations of disease status which can be utilized to infer disease-specific GCNs.

Our finding that marginal improvement in network reconstruction decreases with continued data aggregation suggests that simply addressing statistical considerations due to sample size may have limitations for improving GCNs. Including orthogonal sources of information such as gene-enhancer associations inferred from Hi-C data [52] and transcription factor binding sites from ChIP-seq data [53], in addition to gene expression quantified by RNA-seq in both bulk and single-cell studies, might result in a more accurate understanding of the shared regulatory architecture between genes. Additionally, experimental protocols such as Perturb-Seq [54], which quantifies the transcriptional changes mediated by genetic manipulations on genes, processes, and states, could provide a new avenue for network inference and suggest causal mechanisms and edge directionality.

## Conclusions

We demonstrated that data aggregation improved the inference of consensus and context-specific networks, particularly when properly accounting for latent confounding and between-study variability. While consensus networks prioritized ubiquitous biological processes, context-specific networks captured tissue-specific gene interactions. Further, context-specific networks prioritized variants that are enriched for trait heritability independent of overlap with baseline functional genomic categories suggesting that improving the detection of context-specific gene-gene interactions can shed light on the mechanisms that relate genetic variants to traits. Thus, meaningful data aggregation improves GCN inference and identifies how genes interact to produce complex phenotypes.

## Methods

### A. Data pre-processing and quality control

We downloaded uniformly processed RNA-seq samples from humans using the *recount3* R package [16] and selected 1747 projects that included 30 or more samples each **(Fig. 1A)**. Before normalization, we excluded samples with zero expression across all genes and genes that had zero expression across all samples in a project. We used in-built functions from the *recount3* package to compute the RPKM transformed count matrix, selected genes that were protein-coding, autosomal, and unambiguously mapped to the reference genome [55, 56], and generated the *log*_2_ (*RPKM* + 1) count matrix for each project. Following preliminary processing, we applied a unique data processing pipeline based on the study of origin to exclude samples from micro-RNA and scRNA-seq experiments and summarized replicates. For projects belonging to GTEx, we excluded duplicates (which are labeled as STUDY_NA in the project attribute), as well as samples derived from the chronic myelogenous leukemia (CML) cell line and grouped samples by the tissue of origin to obtain 50 groups and 18828 samples.

Before identifying replicates, we excluded 67 TCGA samples that did not have sample type specified, followed by 39 samples which did not have patient ID present. We then identified replicates in the remaining TCGA samples as those samples that agreed on both patient ID as well as sample type and aggregated by computing the median across all replicates. We then grouped samples by their respective cancer code to obtain 33 groups and 11091 samples.

Finally, for projects from SRA, we began by excluding samples obtained by size fractionation. We then identified replicates, as those samples with an identical experiment accession number and aggregated across replicates by computing the median gene expression, this results in 200499 samples. To exclude microRNA projects that have erroneously been labeled bulk-RNA sequencing experiments, for each sample, we computed the fraction of genes with zero expression and excluded 89,101 samples where this fraction is greater than 50%. The *recount3* database includes manually curated tissue types and experiment types for 30473 samples of 373 SRA studies. The experiment types found included 4 categories, bulk RNA-sequencing, single-cell RNA-seq, small/micro RNA-seq and others. Based on the differences in the sparsity patterns of bulk RNA sequencing and other sequencing modalities, the authors developed a predictor to classify samples as either bulk or single-cell-based. We utilized these predicted labels to restrict our analysis to only bulk RNA-sequencing studies. Since predicted labels were not present for all samples, we excluded samples with keywords such as “single cell”, “scRNA”, “snRNA”, and “single nucleus” in the study abstract if it was found. Further, when the library selection information was available, we restricted our analysis to samples that contained either cDNA or RT-PCR in the library selection field. We also excluded studies which are known scRNA-seq experiments including, SRP096986, SRP135684, SRP166966, SRP200058, and SRP063998. For the remaining studies, we performed a text-based analysis to obtain the Study, Tissue, Organ, Biopsy, Cell, Disease, Source and Description from the metadata `sample_description` field. We then manually annotated 10179 unique combinations of these fields to obtain tissue, cancer status, and disease type. In this manner, we were able to obtain labels for 65361 samples which we grouped on the basis of the study accession IDs to form 884 SRA studies **(Additional File 19: Supplementary File 4)**. While we grouped GTEx samples by tissue of origin, SRA samples by study accession ID, and TCGA samples by cancer accession code, we refer to an individual group of samples as a “study” to simplify nomenclature. Further, we did not distinguish on the basis of disease state or cancer status while organizing the data, until we proceeded to compute the inputs to network inference.

### B. Identifying tissue type and cancer status

Wilks et. al [16] demonstrated that for bulk RNA-seq data from humans, the largest source of variation in gene expression was correlated with tissue or cell type of origin, with the clear clustering of samples belonging to the same tissue extending to the top 4 principal components. We manually refined annotated labels and grouped samples by context using t-SNE dimensionality reduction and clustering 95484 samples and 5999 genes which had non-zero variance across all samples. Since it was not possible to differentiate cancerous and non-cancerous samples using differential clustering in the t-SNE space, we utilized manually annotated labels to restrict the analysis to non-cancerous samples.

Further, to facilitate the greatest possible increase in sample size, we combined similar tissues to form tissue categories **(Additional File 1: Supp. Table I)**. Since differences in gene expression across samples can be driven by factors other than the tissue of origin, such as experimental batch and library size, we obtained the relative enrichment of 64 distinct stromal and immune cell types by comparing the gene expression profile of an admixture sample from bulk tissue to curated gene expression signatures provided by xCell [22]. Once we computed the cell-type enrichment vectors for each sample, we performed t-SNE dimensionality reduction for visualization and clustering.

For 24 tissue categories that have more than 500 samples, we refined our sample selection by eliminating outliers using 6 different methods of outlier calling. These included

1. Z-score-based outlier calling: We computed the z-scores for each reduced dimension independently, samples which have a z-score greater than or equal to 3 in either dimension are labeled as outliers.
2. MAD-based outlier calling: For each dimension we computed the median absolute deviation as given by Eqn. 1

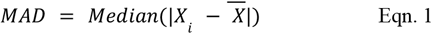

Where 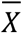, is the mean of all observations. We computed the outlier score in each dimension using Eqn. 2

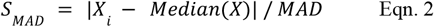

We labelled a sample an outlier if *S* _*MAD*_ is greater than or equal to 3 in either dimension.
3. Tukey outlier calling: We computed the quartiles of each dimension and the *IQR* = *Q*_3_ − *Q*_1_. Samples which have a value lower than *Q*_1_ − 1. 5 × *IQR* or greater than *Q*_3_ + 1. 5 × *IQR* in either dimension are labeled outliers. For each method below, we computed the Mahalanobis distance (MD) for each sample by estimating the covariance matrix *C* across all samples. The MD is given by Eqn. 3, To identify outliers, we used the additional criteria based on the MD outlined below,

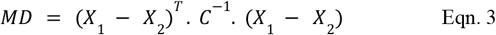
4. Mahalanobis Z-score: Computed the z-scores across the MD metrics and labelled samples with a z-score greater than or equal to 3 as outliers.
5. Mahalanobis MAD: Computed the MAD-based outlier score on the MD using Eqn. 2, And labelled samples with a score greater than or equal to 3 as outliers.
6. Mahalanobis Tukey: Compute the quartiles of the MD estimates and the inter-quartile range (IQR). Label samples with an MD estimate lower than *Q*_1_ − 1. 5 × *IQR* or greater than *Q*_3_ + 1. 5 × *IQR* as outliers.

If a sample within a tissue category is labeled an outlier by at least 2 methods of outlier calling, then we excluded the sample for downstream network reconstruction. While this works with simpler tissue categories such as blood, skeletal muscle or colon where the samples have relatively homogenous cell type compositions, for the immune system, which comprises samples from the distinct myeloid, lymphoid and innate immune systems, these outlier metrics failed due to large intersample variation. Therefore for immune cell types, we first performed K-means clustering with 3 centroids. We annotated the three resulting clusters based on the representation of manually annotated labels in each cluster as either B-cells, myeloid cells (including monocytes and macrophages) and PBMCs w/ T cells. For each cluster, we performed outlier detection using all six methods outlined above and excluded samples that were found to be outliers in any two methods.

### C. Data correction and aggregation

Principal component (PC) based correction methods can account for technical and biological artifacts that confound gene expression measurements and reduce false positives in gene network inference [24]. However, these methods have been applied to one experiment and not across multiple experiments from disparate sources. We systematically compared 4 strategies of data aggregation including,

1. Aggregating data: Identify genes with non-zero expression and non-zero variance across all studies of interest, followed by aggregating the data, quantile normalization, scaling, and PC estimation. We determined the number of PCs to regress using the permutation method described by Buja et. al [25] and Leek et. al [23]. We then obtained the residuals by fitting a linear regression using the number of identified PCs. Finally, we quantile normalized the residuals and scaled each gene to have a mean of zero and unit variance and estimated the empirical covariance matrix.
2. Aggregating data adjusted for confounding: We considered each study individually, quantile normalized, scaled the data, and estimated PCs, followed by the determination of the optimal number of components to be regressed. We then obtained the residual gene expression using a linear model to regress latent variables corresponding to confounding. We aggregated across each individually corrected study by selecting genes that have non-zero expression and non-zero variance across all samples. Finally, we quantile normalized and scaled the aggregated corrected gene expression and estimated the empirical covariance matrix.
3. Unweighted aggregation of covariance matrices: For each individual study, we quantile normalized and scaled the data, followed by PC-based data correction and computed residuals. To the residuals, we applied quantile normalization and scaling and estimated the empirical covariance matrix *S*_*k*_ for each study *k*. Assuming equal likelihood of error from each study we computed the unweighted average of covariance matrices *C*_*unweighted*_ as described in Eqn. 4

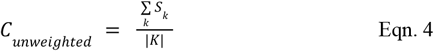

where |*K*| is the total number of studies aggregated.
4. Weighted aggregation of covariance matrices: We repeated the process described previously in the computation of unweighted aggregation of covariance matrices to estimate study-level empirical covariance matrices *S*_*k*_ . We then assumed that studies with a larger sample size provide a better estimate of individual covariances and computed the weighted covariance as in Eqn. 5 And Eqn. 6

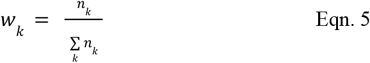

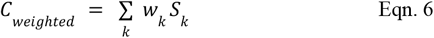

### D. Network reconstruction with graphical lasso

Following the computation of aggregate covariance matrices using the strategies outlined in the previous section, we infer gene regulatory relationships using graphical lasso [13]. The desired network structure is obtained by identifying the precision matrix, *Θ* = *Σ*^−1^ that maximizes the penalized log-likelihood given by Eqn. 7, where *C* is the estimated covariance matrix.

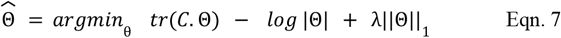

We estimated the precision matrix *Θ* across a range of *λ* between 0.04 to 1.00 in intervals of 0.02. For genes *p* and *q*, an edge connecting them exists if the corresponding entry in the precision matrix is non-zero as given by Eqn. 8

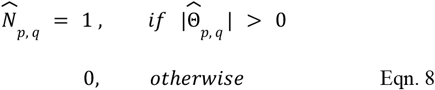

### E. Network evaluation to determine the optimal aggregation strategy

We applied all 4 methods of data aggregation to infer networks from non-cancerous samples using two data partitions - GTEx and SRA non-cancer. Since GTEx did not include any cancerous samples, we organized the samples by tissue-type and excluded any tissues with fewer than 15 samples, in this case, Kidney-Medulla. For the SRA non-cancer, we select samples which were annotated to be non-cancerous and organize them by the study accession IDs, we then exclude any studies which had fewer than 15 samples, resulting in 566 selected studies. For each data partition, we evaluated network improvement with sequential data aggregation as follows,

1. SRA non-cancer: An aggregate network inferred across 566 SRA studies from varying tissues that are non-cancerous. We order the studies by increasing sample size and aggregate 1, 100, 200, 300, 400, 500 and 566 studies to estimate empirical covariance matrices and infer networks using graphical lasso across *λ* ranging from 0.04 to 1.00.
2. GTEx non-cancer: An aggregate network inferred across 49 GTEx tissues and cell lines. We order tissues in increasing order of sample size and aggregate tissues in increments of 10 to obtain networks corresponding to 1, 10, 20, 30, 40 and 49 tissues across a range of *λ* between 0.04 and 1.00.

Details of the number of studies, samples, median sample size and principle components regressed are provided in **Additional File 3: Supp. Table II**. Following network inference, we evaluated the improvement in network inference as a function of sample size and method of data aggregation using two metrics, held-out log-likelihood and enrichment of known biological pathways. The metrics are computed as follows,

1. Held-out log-likelihood: For networks inferred using SRA data, the test set consisted of 6 GTEx tissues including Adipose Subcutaneous, Cells cultured fibroblasts, Heart left ventricle, Artery aorta, and Colon sigmoid. To evaluate networks inferred using GTEx samples we selected SRA studies with matched sample sizes including SRP116272, SRP151763, SRP150552, SRP174638, and SRP187978. For each test study, we quantile normalized and scaled the data, followed by PC-based data correction and computed residuals. We then quantile normalized and scaled the residuals and computed the corresponding covariances matrices *S*_*i*_ for the *i*^*th*^ study. We computed the log-likelihood of the data observed in each held-out study as shown in Eqn. 9

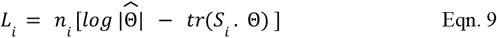

We then computed the average log-likelihood across all test data sets *i ∈ I* as given by Eqn. 10 where |*I*| is the total number of test data sets.

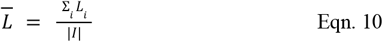
2. Prediction of known biological pathways: We downloaded pathway information from KEGG, Biocarta and Pathway Interaction Database from Enrichr and selected those pathways that were annotated as canonical pathways by MSigDB. **Additional File 4: Supp. Table III** provides details on the number of gene sets from each source. Any pair of genes that have at least one pathway in common were assumed to have a true functional relationship. In total, we identified 9648013 such edges (*U*). For each inferred network 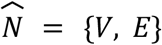, where *V* is the set of genes over which we perform network reconstruction, and *E* is the set of edges between the gene pairs, we restricted the set of universal pathway edges to those between genes *G*_1_, *G*_2_ *∈ V*. If *S*_*all*_ is the set of edges between all possible gene pairs such that 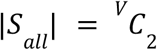, then *U*^*C*^ = *S*_*all*_ *\ U*, and *E*^*C*^ = *S* _*all*_ *\ E*, we constructed the contingency table as follows,
  a. True positives (TP) = |*U ∩ E*|
  b. False positives (FP) = |*U*^*C*^ *∩ E*|
  c. True negatives (TN) = |*U*^*C*^ |*U ∩ E*^*C*^ |
  d. False negatives (FN) = |*U*^*C*^*∩ E*^*C*^ |

We computed the precision, recall, and F1-score as described by Eqn. 11, Eqn. 12, Eqn. 13.

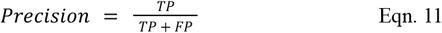

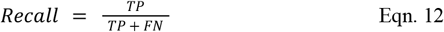

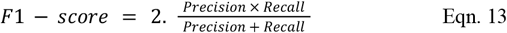

## F. Inference of consensus and tissue context-specific networks

We inferred consensus networks across samples from disparate tissues and cell types to capture shared biological pathways across contexts. Since weighted covariance aggregation yielded the best results across the 4 methods of data correction and aggregation, we first grouped SRA samples by the study accession ID, GTEx samples by the tissue of origin, and TCGA samples by the cancer code. To estimate the universal consensus network we did not exclude any samples. Thus, we aggregated 65361 samples from 884 SRA studies with a median sample size of 44, 11091 samples from 33 TCGA studies with a median sample size of 309, and 18824 GTEx samples from 49 tissues and a median sample size of 286. Prior to estimating non-cancer consensus networks, we first exclude all cancerous samples from each study, and only retain studies with 15 or more samples. Similarly, prior to estimating cancer consensus networks, we only include cancerous samples, and exclude studies with fewer than 15 samples. Since weighted covariance aggregation yielded the best results across the 4 methods of data correction and aggregation, we first etimated and adjusted for unknown technical confounders using Principal component analysis (PCA) in each group of samples. We computed the weighted empirical covariance matrix as detailed in Section II.c. We then inferred the network structure by optimizing penalized log-likelihood Eqn. 7 And thresholded the entries in the precision matrix as described by Eqn. 8 for a range of penalization parameters *λ* ranging from 0.08 - 1.00.

We inferred context-specific networks in 27 contexts with 500 or more samples. Details of the number of samples, studies, and median sample size across studies for each context are provided in **Additional File 2: Supp. Fig 13**. For 20 contexts including adipose, B cells, blood, breast, cardiac, central nervous system, colon, esophagus, fibroblasts, intestine, kidney, liver, lung, nervous system, pancreas, prostate, skeletal muscle, skin, stomach, and vascular we inferred networks either using GTEx sample only or by aggregating context-specific samples from recount3 which included GTEx. For each context-specific network, we first performed PC-based data correction within each study followed by covariance estimation and aggregation by weighting the covariance matrix with the proportion of study-specific sample size to the total number of samples as detailed in Eqn. 5 and Eqn. 6. We then inferred GCNs using graphical lasso using Eqn. 7 and Eqn. 8

## G. Evaluating the impact of data aggregation on the inference of consensus and context-specific networks

To evaluate the improvement in the performance of consensus networks with data aggregation, we computed the precision, recall, and F1-score of observing known gene co-regulatory pairs compiled from KEGG, Biocarta and Pathway Interaction Database which were annotated as canonical pathways by MSigDB as detailed in **Methods (Section E)**. Specifically, we compared the F1-scores corresponding to the universal, non-cancer, and cancer consensus networks for network densities between 5000 and 500,000 edges.

In order to evaluate the impact of data aggregation on context-specific networks, we selected two contexts with the highest number of samples - blood and CNS. We obtained 64 SRA studies that were annotated to blood, and 40 SRA studies annotated to CNS. We sequentially aggregated SRA studies five at a time in each context in the increasing order of sample size and inferred GCNs using weighted aggregation of covariance matrices and graphical lasso as detailed in **Methods (Sections C, D)** over a range of penalization parameters *λ* varying from 0.10 to 1.00. For each data aggregation setting, we evaluated the network performance over a range of network densities by computing the held-out log-likelihood of GTEx samples belonging to the concordant tissue in a manner analogous to **Methods (Section E)**. Specifically, to evaluate the inferred blood networks, we utilized the empirical covariance matrix computed from 850 blood GTEx samples. Since GTEx contained 13 brain regions which were all annotated to the CNS context, we computed the held-out log likelihood corresponding to each brain region and computed the mean across regions to obtain a scalar measure of the network’s generalizability.

Additionally, for 6 contexts, adipose, blood, CNS, liver, lung, and skin we compared networks inferred solely from GTEx samples to GCNs inferred from data aggregation based on the inclusion of known tissue-specific protein-protein interactions (PPIs). First, we selected networks that were inferred with the value of the penalization parameter between 0.12 and 0.50. We then downloaded tissue-specific PPIs present in PPT-Ohmnet_tissues-combined.edgelist.gz from SNAP which is compiled across 144 unique tissues [57–59]. We mapped the tissue labels present in SNAP to our contexts studied as given in **Additional File 20: Supp. Table XV**. After converting the protein identifiers using the R package `org.Hs.eg.db,` we performed Fisher’s test to compute the odds ratio of finding tissue-specific PPIs in our context-specific networks when compared to a fully connected network as background. Specifically, we computed the following contingency matrix that we used as input to the R function `fisher.test()`where *E* is the set of edges present in our inferred network, *P*_*t*_ is the set of concordant tissue-specific PPIs, and *E*^*c*^ is the set of all edges absent in the network which is obtained by *U \ E*, where *U* is the set of all possible edges between the nodes, and 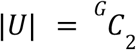 where *G* is the set of genes over which we perform network inference.

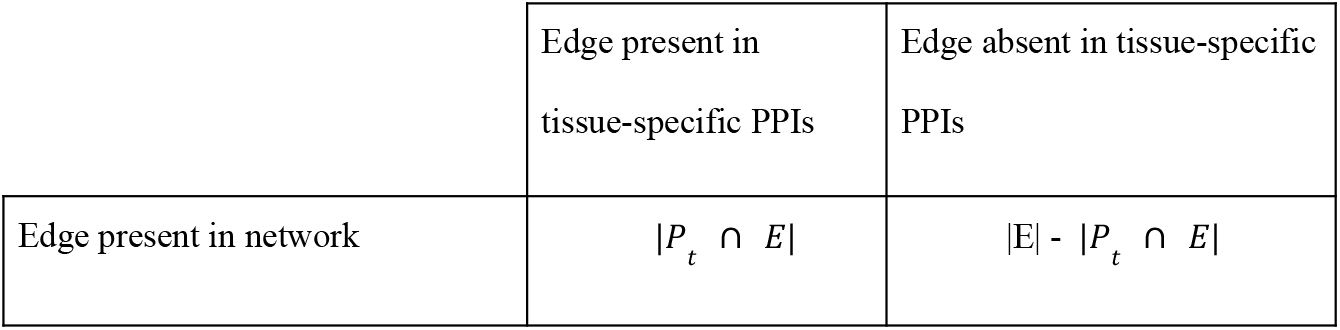
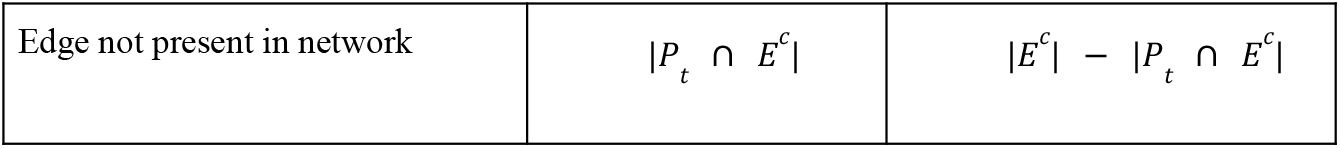

|  | Edge present in tissue-specific PPIs | Edge absent in tissue-specific PPIs |
| --- | --- | --- |
| Edge present in network | $\vert P_t \cap E\vert $ | $\vert E\vert - \vert P_t \cap E\vert $ |
| Edge not present in network | $\vert P_t \cap E^c\vert $ | $\vert E^c\vert - \vert P_t \cap E^c\vert $ |

To compare between context-specific networks inferred solely from GTEx to those inferred across aggregated data, we performed the Wilcoxon rank-sum test to compare the odds ratios from each setting.

## H. Sparsity parameter selection for consensus and context-specific networks

Previous analysis of gene regulatory networks indicates that eukaryotic [60] and prokaryotic transcriptional [61] networks exhibit an approximately scale-free degree distribution which concurs with the assumption that few transcription factors have the potential to regulate an multitude of target genes [35]. The power law degree distribution follows Eqn. 14 where *p*(*k*) is the probability of a random node having *k* connections and *γ* is the power law scaling exponent.

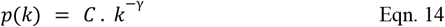

Taking the logarithm, we get,

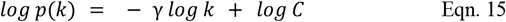

For each of the 3 consensus networks and 27 tissue-context specific networks, we used the R package `igraph` to estimate the degree distribution for each value of *λ* and the function `lm()`to estimate the slope (− *γ*) and *R*^2^ . Finally, we computed the 95% confidence interval (CI) for the slope using the R function `confint().` We then selected networks such that 2 *≤ LL*(*γ*) *≤ UL*(*γ*) *≤* 3, and the *R*^2^ *> 0. 8* where *LL*(*γ*) and *UL*(*γ*) correspond to the lower and upper limits of the 95% CI of *γ*. Details pertaining to the selected consensus networks and their corresponding values of the penalization parameter *λ*, network densities, regression slope and *R*^2^ are provided in **Additional File 5: Supp. Table IV**. Similarly, the selected network densities, values of the regression slope and *R*^2^ are provided for context-specific networks inferred from GTEx only and across all aggregated samples in 6 contexts which were used in downstream analysis is in **Additional File 6: Supp. Table V**.

## I. Computing gene centrality measures based on network structure

Using measures of network connectivity, we computed centrality scores for each gene in the network. Given a weighted undirected graph *G*, first, we normalized the edge weights If *E* is the list of all edges in the network (excluding diagonals) and *E*_*p, q*_ is the weight of an edge connecting genes *p* and *q*. Then the normalized weight *E* _*p, q*_ = *E*_*p, q*_ */ max*(*E*_*i, j*_) for all *i, j ∈ Nodes*(*G*) such that *i ≠ j*. Using normalized edge weights we computed the following centrality scores for each gene *p*,

a. Betweeness(*p*): The betweeness centrality captures the number of shortest paths in a network that passes through the gene *p*. The shortest path between nodes *i, j ∈ Nodes*(*G*) is the path where the sum of the edge weights is minimum.
b. Closeness(*p*): The closeness centrality captures the distance between a gene *p* to all other network nodes. We first obtained the weighted distance between gene *p* and gene *i* as given by Eqn. 18

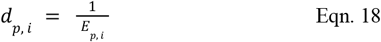

If the nodes *p* and *i’* are disconnected then we set *d*_*p, i’*_ to 0. Finally, the closeness centrality of a gene *p* is given by Eqn. 19

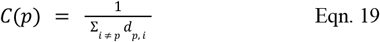
c. Degree(*p*): The degree of a gene *p* corresponds to the number of neighbors connected to *p*.
d. Maximum weight(*p*): The maximum weight of a node is the maximum weight of the edges that are connected to the gene *p*.
e. Page rank(*p*): The page rank of a node *p* is defined by Eqn. 20,

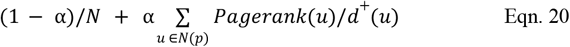 Where *α* is a damping factor, *N*(*v*) are the in-neighbors of node *p*, and *d*^+^(*u*)is the out-degree of *u*. Because *G* is an undirected graph, pagerank treats it as a directed graph by making edges bi-directional. Specifically, in addition to the number of neighbors that a node *p* has, page rank incorporates information pertaining to the connectivity of the neighboring nodes.
f. Strength(*p*): The strength of a node is defined as the sum of the weights of all edges connected to the gene *p*.

## J. Enrichment of specific pathways among central genes in consensus and context-specific networks

First we developed a general method to test for the enrichment of genes annotated to a specific GO term among network central genes. For a given network *N* = *{V, E}* where *V* is the set of nodes and *E* the set of edges, we estimated the degree of each node as described in **Methods (Section I)** and estimated the maximum degree across all nodes. We then determined a series of degree thresholds, starting at 5 to the maximum degree in increments of 5. For a particular degree threshold, *D*_*t*_ we obtained all the network nodes with a degree *D*_*i*_ *≤ D*_*t*_, which forms the test set of genes *T*, such that *T ⊆ V*. To determine the enrichment of genes associated with a particular GO term corresponding to a biological process (*GOG*), we first computed the following contingency matrix where *G* is the set of 19950 protein-coding genes where *T*^*c*^ = *G\T*

|  | Gene present in GO term | Gene absent in GO term |
| --- | --- | --- |
| Gene present in test set | $\vert T \cap GoG\vert $ | $\vert T\vert - \vert T \cap GoG\vert $ |
| Gene absent in test set | $\vert T^c \cap GoG\vert $ | $\vert T^c\vert - \vert T^c \cap GoG\vert $ |

We then performed Fisher’s test with the alternative hypothesis that the odds ratio of finding GO related terms is greater in the test set *T* and estimated p-values.

To evaluate the universal, non-cancer and cancer consensus networks, we first selected the value of the penalization parameter such that the resulting network density was ∼ 7000 edges. We then determined the odds ratio of finding genes belonging to ubiquitous biological processes among network nodes selected by successively increasing degree thresholds. Further, we computed the odds ratio of finding genes belonging to ubiquitous processes in blood and CNS networks inferred across all samples from *recount3* with ∼ 7000 edges.

To evaluate a context-specific network such as blood, we first selected a blood network inferred solely from GTEx samples and a network inferred by data aggregation across all samples from *recount3* such that the resulting network density was ∼7000, and the node degree distribution was scale-free. As negative controls, we included a different context-specific network inferred across aggregated samples, the universal and non-cancer consensus network. For blood, we specifically tested for the enrichment of processes such as leukocyte migration, leukocyte activation and blood coagulation. We repeated this analysis for 5 other context-specific networks - CNS, skin, lung, liver, and adipose by selecting related GO terms and determining the enrichment of functional genes among network central genes. A complete list of GO terms that were tested for each context is described in **Additional File 7: Supp. Table VI**.

## K. Excess overlap of genes grouped by centrality measures in consensus with known evolutionarily constrained and functionally prioritized gene sets

We group genes for each consensus network such that the first bin includes genes with no edges (with a degree of zero), the second bin contains nodes with a degree in the lowest 25^th^ percentile, the third bin includes nodes with a degree in the 25^th^ - 50^th^ percentile, the fourth bin includes genes with degree in the 50^th^ - 75^th^ percentile and the fifth and final bin includes genes with a degree greater than the 75^th^ percentile. For genes in a given bin, we compute the excess overlap with known 9 evolutionarily constrained and functional gene sets used by S.Kim et al. which include,

1. High pLI genes [62]: 3,104 loss-of-function (LoF) genes with pLI > 0.9, i.e., strongly depleted for protein-truncating variants.
2. High S_het_ genes [63]: 2,853 constrained genes with S_het_ > 0.1 i.e., strong selection against protein-truncating variants.
3. High missense Z-score [64]: 1,440 constrained genes strongly depleted for missense mutations, with exp_syn *≥* 5, syn_z_sign < 3.09, and mis_z_sign > 3.09, as retrieved in Lek et al [62].
4. High Phi [65]: 588 LoF-constrained genes with a probability of haploinsufficiency (Phi) > 0.95.
5. MGI essential genes [66–68]: 2371 genes for which a homozygous knockout resulted in pre-, peri-, post-natal lethality
6. eQTL deficient [69]: 604 genes with no significant variant-gene association in all 48 tissues in GTEx v.7 single-tissue cis-eQTL data.
7. OMIM [70]: 2,266 genes deposited in the Online Mendelian Inheritance in Man (OMIM), as retrieved in Petrovski et al [71].
8. ClinVar [72]: 5,428 genes with a pathogenic or likely pathogenic variant with no conflict among studies.
9. DrugBank [73]: 373 genes whose protein products are human targets of FDA-approved drugs with known mechanisms of action.

The complete list of genes belonging to each of these categories is provided in **Additional File 8: Supp. Table VII**. We computed the expected and standard error of excess overlap between genes in each bin *G*^*i*^_*b*_ for *i* = 1, 2, 3, 4, 5 and the reference gene set *j, G* ^*j*^_*r*_ using Eqn. 21 - 24, where *G*_*tOt*_ represents all network nodes.

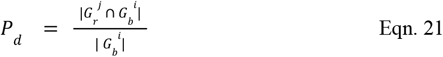

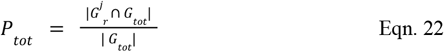

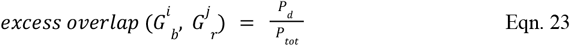

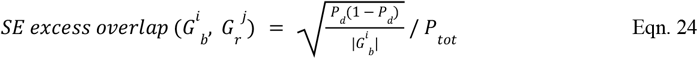

## L. Identifying biological processes associated with shared and distinct hub genes from non-cancer and cancer consensus network

In accordance with the scale-free criterion we selected the non-cancer network inferred using a penalization parameter λ = 0.18 resulting in a network density of 7355 edges and the cancer consensus network corresponding to the penalization parameter λ = 0.24 and a network density of 7552 edges to compare the biological processes which are represented by shared and distinct network hubs. First, we defined hub genes as network nodes with a closeness centrality in the 90th percentile independently for each consensus network, such that the non-cancer hub genes are given by *H* _*N*_ and the cancer hub genes are given by *H*_*C*_ . We then identified hub genes shared between cancer and non-cancer consensus networks *H*_*S*_ = *H*_*N*_ *∩ H*_*C*_, non-cancer specific hub genes *H*_*NS*_ = *H*_*N*_ *∩ H*_*C*_ *‘*, and cancer-specific hub genes as *H*_*CS*_ = *H*_*C*_ *∩ H*_*N*_ *‘* (**Additional File 10: Supp. Table IX**). We then used the `GOTermFinder` tool [74] to identify GO terms that are shared by the genes in the sets *H*_*S*_, *H*_*NS*_, and *H*_*CS*_ .

## M. Examining the differences in the degree distribution of tissue-specific vs. general transcription factors in consensus and context-specific networks

We selected context-specific networks inferred from CNS, blood, cardiac, skin, lung, skeletal muscle, and pancreas samples from *recount3* which had an approximate density of 7,000 edges (**Additional File 21: Supp. Table XVI**). Additionally, we selected values of the penalization parameter *λ* which yielded universal and non-cancer consensus networks with an approximate density of 7,000 edges. We referred to Pierson et al. [36] to obtain a list of tissue-specific and general transcription factors which are provided in **Additional File 9: Supp. Table VIII**. In order to select a background set of non-TFs, we first obtained the intersection of genes which were included in each network considered, then we excluded both general and tissue-specific TFs from this list, and randomly select 100 of these genes. The selected background is provided in **Additional File 9: Supp. Table VIII**. For each network, we compared the degree distribution of tissue-specific and general transcription factors to non-TFs and the degree distribution of tissue-specific to general transcription factors using the Wilcoxon rank-sum test. Across all tests, we adjusted for multiple hypothesis correction using the Holm-Bonferroni method.

## N. Non-negative matrix factorization to determine shared co-regulatory relationships in similar tissues

We selected context-specific networks with a density of ∼7000 across 27 contexts for which we inferred GCNs by aggregating samples assigned to the context from *recount3* (**Additional File 21: Supp. Table XVI**). Further, the selected networks were in accordance with the scale-free selection criterion detailed in **Methods (Section H)**. For each network, we identified hub genes as network nodes with a degree centrality in the 95th percentile. Across the 27 contexts, we found 3682 unique hubs. We subset to hubs that are present in at least 2 contexts which results in 1956 hubs. We then used the R package `RcppML` to perform non-negative matrix factorization to learn 8 underlying factors to group similar patterns of hub genes. Specifically, *H* is a binary matrix of dimensions *N*_*Hubs*_ × *N*_*c*_, where *N*_*c*_ is 27 the number of contexts. *H*_*i, j*_ = 1 when the *i* ^*th*^ gene is a hub gene in context *j* and *H*_*i, j*_ = *0* otherwise. We then obtain matrices *W* of dimensions *N*_*Hubs*_ × *8*, and *C* of dimensions *N*_*c*_ × *8* by solving the following optimization function,

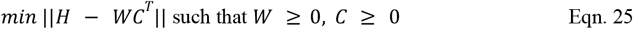

Finally, we chose a solution after 10 iterations that resulted in the greatest sparsity, i.e. smallest values of ||*W*||_1_ and ||*C*||_1_ . To interpret the resulting context cluster, we examine the genes that contributed the most to the corresponding factor. Specifically, for a given factor, *W[:, p]*, we first obtain genes with loadings in the 80% percentile. Then for each gene, we compute the maximum difference between the loading of the gene on factor *p* to all other factors and select genes where this difference is *≥* 5e-5. Thus we obtain a set of factor-specific hub genes *G*_*p*_ (**Additional File 12: Supp. Table X**). We then used the `GOTermFinder` tool [74] to identify GO terms that were more likely to be present in the set *G*_*p*_ than a background of all protein-coding genes by estimating the odds ratio and p-value by applying the hypergeometric test.

## O. Stratified LD-Score regression to quantify the heritability enrichment of variants proximal to central network genes in consensus and context-specific networks

We applied stratified LD-score regression (s-LDSC) [21] to quantify the contribution of node centrality estimated from our consensus and context-specific GCNs towards explaining complex-trait heritability. First, we transformed all centrality scores derived from our universal consensus, non-cancer consensus, blood and CNS context-specific networks to lie between 0 and 1. Next, we annotated SNPs within 100 Kb of a gene with the centrality score assigned to the gene. If an SNP was within 100 Kb of more than one gene, we assigned the maximum centrality score across all corresponding genes to the SNP. We generated six network centrality-based annotations **(Methods, Section I)** corresponding to each network studied and estimated their heritability enrichment and the standardized effect size (τ *_*c*_) of an annotation *c* as described by S. Kim et. al [40].

Given an SNP *j* with a trait-specific effect size of *β*_*j*_ and a corresponding annotation value *a*_*c,j*_ pertaining to category *c*, the variance of the effect size *β*_*j*_ is assumed to be a linear additive combination of the annotation *c* as given by Eqn. 25.

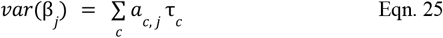

Here τ_*c*_ is the per-SNP contribution of the category *c* to the heritablity of the trait.

s-LDSC then estimates τ_*c*_ using the regression model specified by Eqn. 26.

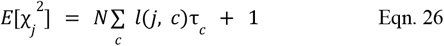

Where N is the number of samples in the GWAS and *l*(*j, c*) is the LD score of SNP *j* to the annotation *c* computed as given by Eqn. 27.

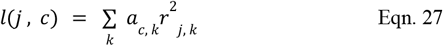

Where *a*_*c, k*_ is the annotation value of SNP *k* and *r*^2^ _*j, k*_ is the correlation between SNPs *j* and *k*. We can then compute the enrichment of an annotation as the proportion of heritability explained by SNPs in a given annotation divided by the proportion of SNPs in the annotation. This definition can be extended to continuous annotations as given by Eqn. 28.

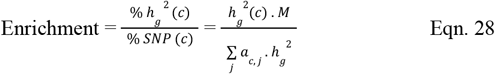

Where *h g*^2^ (*c*) is the heritability captured by the *c* ^*th*^ annotation, *h* _*g*_^2^ is the estimated SNP-heritability and *M* is the total number of SNPs used to compute *h*_*g*_ ^2^ which in this case is 5,961,159. When the annotation is enriched for trait heritability, the computed enrichment is > 1. To compute the significance of the enrichment, we used the block jack-knife method presented in previous studies [21, 40, 75, 76].

Finally, we computed the standardized effect size τ_*c*_ * i.e. the proportionate change in the per-SNP heritability associated with a single standard deviation increase in the value of the annotation *c* conditioned on all other annotations in the model as given by Eqn. 29.

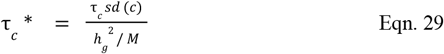

Where *sd*(*c*) is the standard deviation of the annotation *c*. τ_*c*_ * thus captures the effect unique to the focal annotation, *c* unlike enrichment. The significance for the effect size of each annotation is computed by assuming τ_*c*_ * */ se*(τ_*c*_ *) *∼ N*(*0*, 1) as done previously [40, 75, 77].

We used European samples from the 1000G phase 3 project [49] to obtain reference SNPs. Regression SNPs were obtained from HapMap 3 [78] and regression weights were obtained excluding SNPs in the major histocompatibility complex (MHC) from 1000G_Phase3_weights_hm3_no_MHC.tgz . We estimated heritability enrichment and annotation effect size τ _*c*_*, using two model settings. In the first, we included annotations corresponding to variants located in a 100 Kb window of all protein-coding genes (all-genes annotation), in the second setting we included annotations corresponding to variants located in a 100 Kb window of all protein-coding genes and 97 annotations present in the baseline-LD model v2.2 (all-genes + baseline). Specifically, we downloaded the 1000G_Phase3_baselineLD_v2.2_ldscores.tgz file which included 97 variant annotations such as the annotations provided by Gazal et. al [79], Hujoel et. al [80], promoter and enhancer specific annotations from Villar et. al [81], and promoter and enhancer age annotations from Marnetto et. al [82].

To examine the heritability enrichment and effect size of annotations derived from the universal and non-cancer consensus networks which were selected based on the scale-free distribution of node degree detailed in **Methods (Section H)** we used two sets of traits.

1. 42 traits considered by S. Kim et al, such that the Z-score of total SNP-heritability was at least 6 and the genetic correlation between any pair of traits estimated by cross-trait LDSC was lower than 0.9. A complete list of traits is provided in **Additional File 15: Supp. Table XI**
2. 219 UKBB traits with a heritability Z-score >= 7, and the genetic correlation estimated by cross-trait LDSC is lower than 0.5. A complete list of traits is provided in **Additional File 16: Supp. Table XII**.

To examine the heritability enrichment of blood context-specific networks derived from samples solely from GTEx and inferred across all aggregated samples we utilized GWAS summary statistics corresponding to 9 blood-derived traits including, ulcerative colitis [83], rheumatoid arthritis [84], and Crohn’s disease [83], as well as eczema, white cell count, red cell count, red blood cell width, platelet count, eosinophil count from UKBB (**Additional File 17: Supp. Table XII**). Networks derived from blood samples were selected based on the scale-free criterion for node degree distribution detailed in **Methods (Section H)**.

To examine the heritability of CNS context-specific networks derived from samples solely from GTEx and inferred across all aggregated samples we utilized GWAS summary statistics from the Psychiatric Genomics Consortium [85] corresponding to Alzheimer’s disease [86], epilepsy [87], Parkinson’s, bipolar disorder [88], smoking cessation [89], schizophrenia [90], major depressive disorder [91], number of drinks per week [89], and waist-hip-ratio adjusted BMI from [UKBB](https://nealelab.github.io/UKBB_ldsc) (**Additional File 18: Supp. Table XIV**). Networks derived from CNS samples were selected based on the scale-free criterion for node degree distribution detailed in **Methods (Section H)**. Following the computation of heritability enrichment and τ _*c*_* for each trait, as well as their respective standard deviations we performed a random-effects meta-analysis across traits by using the function `meta.summaries()` from the R package `rmeta`.

## Supporting information

Supplemental Table III

Supplemental Table VII

Supplemental Table VIII

Supplemental Table XII

Supplemental Table XIII

Supplemental Table XIV

Supplemental Table XVI

## Ethics Declarations

### Ethics approval and consent to participate

Ethical approval is not applicable for this manuscript.

### Consent for publication

Consent for publication is not applicable for this manuscript.

### Availability of data and materials

Scripts to reproduce the analysis and figures included in this manuscript can be found at: https://github.com/prashanthi-ravichandran/recount3_networks

Networks and annotations used in this project can be obtained from: **10.5281/zenodo.10480999**

### Authors’ contributions

P.R., P.P.., and A.B. conceived the project. P.R., P.P., and A.B. designed the analyses. P.R. and P.P. performed the analyses. R.K. and K.H. contributed feedback to experimental design and interpretation. P.R., R.K., and A.B. organized and wrote the paper with input from all authors.

## Acknowledgements

The authors would like to thank helpful discussions from Battle lab members throughout the course of this work, particularly Joshua Weinstock and Eric Kernfeld for code review and manuscript feedback.

## Supplementary Information

**Additional File 1:** Supp. Table I

Assignment of tissue labels obtained by manual curation to 48 contexts.

**Additional File 2:** Supp. Fig 1 - 19

**Supp. Fig 1**. Identification of outlier samples across contexts. **Supp. Fig 2**. Immune cells were separated into three clusters: Melyoid, PBMCs/T cells, and B cells. **Supp. Fig 3**. Comparison of GTEx to Tissue consensus networks across tissue contexts. **Supp. Fig 4**. Characteristics of PCs at varying levels of data aggregation. **Supp. Fig 5**. ANOVA of top 10 PCs at varying levels of data aggregation. **Supp. Fig 6**. Prevalence of tissues in GTEx and SRA. **Supp. Fig 7**. Held-out Log-likelihood calculated reciprocally for GTEx and SRA. **Supp. Fig 8**. F1 scores calculated reciprocally for GTEx and SRA. **Supp. Fig 9**. Regression estimate (β) and standard errors of the effect of sample size (N) on network density across varying penalization parameters . **Supp. Fig 10**. Held-out log-likelihood of networks inferred by sequential data aggregation of GTEx and SRA studies. **Supp. Fig 11**. Linear model between the log empirical degree distribution and log degree for consensus networks. **Supp. Fig 12**. Odds ratio for ubiquitous biological processes across networks. **Supp. Fig 13**. Odds ratio for tissue specific biological processes across networks. **Supp. Fig 14**. Excess overlap of functional gene sets across networks. **Supp. Fig 15**. Excess overlap of functional gene sets across context-specific networks. **Supp. Fig 16**. S-LDSC results meta-analyzed across 42 traits for annotations from consensus networks. **Supp. Fig 17**. S-LDSC results meta-analyzed across 219 UKBB traits for annotations from consensus networks. **Supp. Fig 18**. S-LDSC results meta-analyzed across 9 blood traits. **Supp. Fig 19**. S-LDSC results meta-analyzed across 9 CNS traits.

**Additional File 3:** Supp. Table II: Details pertaining to the number of samples, median sample size across aggregated studies, number of PCs regressed when data is aggregated prior to correction for technical, and median number of PCs regressed from each study prior to aggregation for sequential aggregation of 566 non-cancerous studies from SRA and 49 studies from GTEx.

**Additional File 4:** Supp. Table III: Specification of data sources used in the compilation of pathways including the number of gene sets obtained from each source.

**Additional File 5:** Supp. Table IV: Selection of penalization parameter Lambda for consensus networks based on the scale-free criterion for degree distribution.

**Additional File 6:** Supp. Table V: Selection of penalization parameter Lambda for context-specific networks based on the scale-free criterion for degree distribution.

**Additional File 7:** Supp. Table VI: Details of GO terms that were tested for enrichment as a function of degree thresholds in consensus and context-specific networks.

**Additional File 8:** Supp. Table VII: Details of evolutionarily constrained and functional gene sets for which excess overlap is computed across network nodes binned by degree centrality.

**Additional File 9:** Supp. Table VIII: List of general and tissue-specific transcription factors derived from Pierson. et. al, as well as randomly selected background genes used to evalute the differences in the centrality scores obtained from consensus and context-specific networks

**Additional File 10:** Supp. Table IX: Hub genes that are shared and distinct between non-cancer consensus network with 11,110 edges (λ = 0.16) and the cancer consensus network with 10,482 edges (λ = 0.22).

**Additional File 11:** Supplementary File 1: Enrichment of GO terms in cancer-specific hub genes.

**Additional File 12:** Supp. Table X: Hub genes selected to describe each factor such that, the loading of the gene on the factor was in the 80th percentile, and the minimum difference between the weight of the gene on the selected factor and all other factors is >= 5e-5.

**Additional File 13:** Supplementary File 2: Enrichment of GO terms among hub genes which are specific to Factor 7 which results in the grouping of PBMCs/ T cells.

**Additional File 14:** Supplementary File 3: Enrichment of GO terms among hub genes which are specific to Factor 6 which results in the grouping of hESCs, iPSCs, and multipotent cells.

**Additional File 15:** Supp. Table XI: 42 Independent traits examined by S.Kim et. al, and their corresponding references, heritability, Z-score and number of individuals in the GWAS

**Additional File 16:** Supp. Table XII: 219 UKBB traits that are selected based on having a Z-score >= 7, and pairwise correlation between traits is lesser than or equal to 0.5 as estimated by LDSC.

**Additional File 17:** Supp. Table XIII: 9 blood-related traits which were used to examine heritability enrichment of blood specific networks.

**Additional File 18:** Supp. Table XIV: 9 CNS-related traits which were used to examine heritability enrichment of CNS specific networks.

**Additional File 19:** Supplementary File 4: Manually curated annotations corresponding to tissue, cancer status and disease type of SRA samples based on text-based analysis of Study, Tissue, Organ, Biopsy, Cell, Disease, Source and Description from the metadata sample_description field.

**Additional File 20:** Supp. Table XV: Mapping between annotation categories present in tissue-specific PPI interactions obtained from SNAP and contexts across which context-specific networks were inferred.

**Additional File 21:** Supp. Table XVI: Selection of context-specific networks used in matrix factorization of shared hubs and edges.

