## Supplementary Figures for "Aggregation of *recount3* RNA-seq data improves inference of consensus and tissue-specific gene co-expression networks"

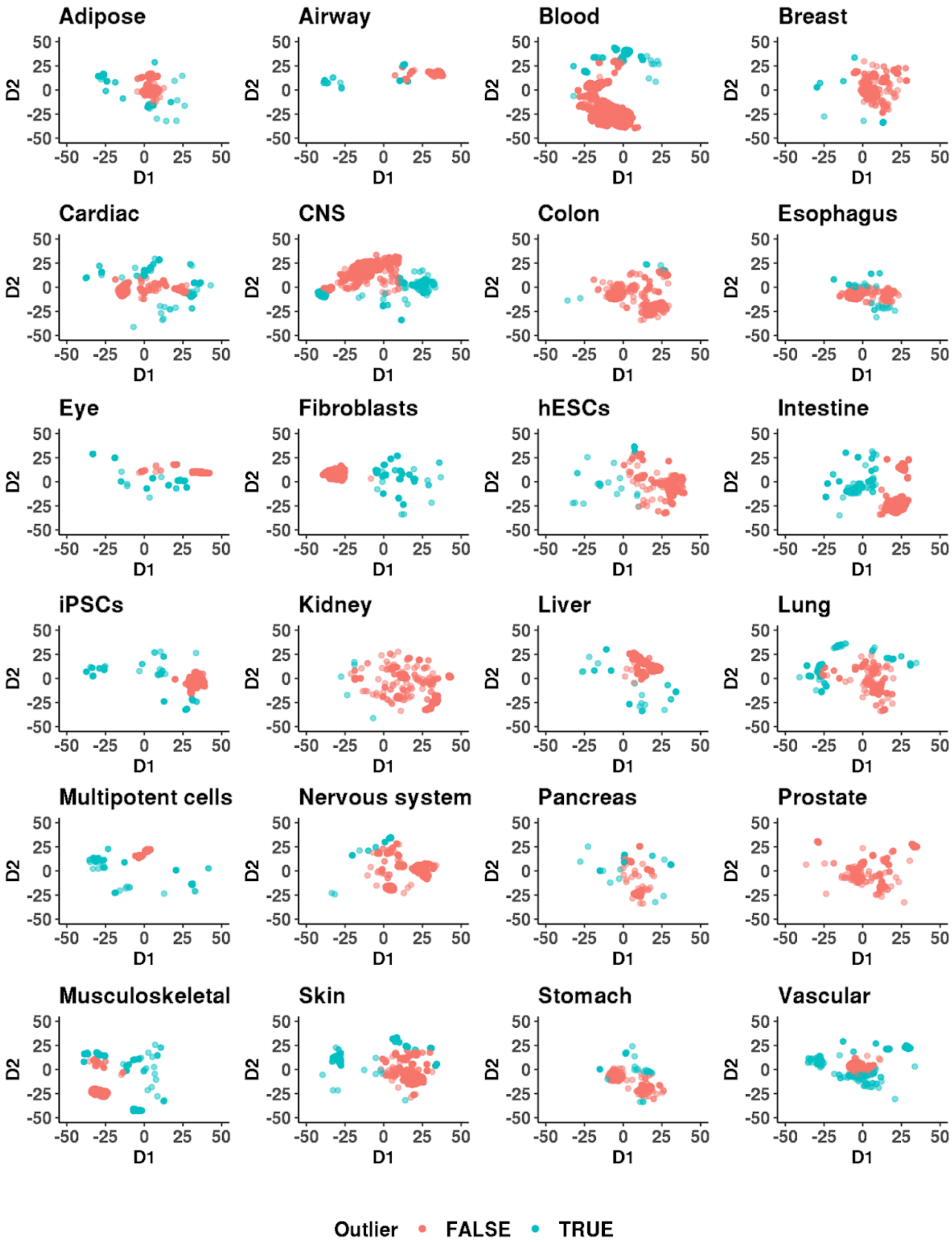

**Supp. Fig 1**

A sample is labelled an outlier if it is identified as an outlier based on either the Z-score, Median Absolute Deviation and Tukey fences on t-SNE projection in each dimension or the Mahalanobis distance computed on the t-SNE projections. Across all tissue contexts, we retained > 67% of samples (93.85% adipose, 76.96% airway, 90.53% blood, 97.04% breast, 89.3% cardiac, 86.16% central nervous system (CNS), 98.48% colon, 97.82% esophagus, 84.65% eye, 84.54% fibroblasts, 92.53% human embryonic stem cells (hESCs), 93.9% intestine, 94.5% induced pluripotent stem cells (iPSCs), 99.63% kidney, 98.12% liver, 82.26% lung, 83.19% multipotent cells, 94.5% nervous system, 79.62% pancreas, 100% prostate, 67.05% musculoskeletal system, 85.12% skin, 94.15% stomach, and 76.74% vascular).

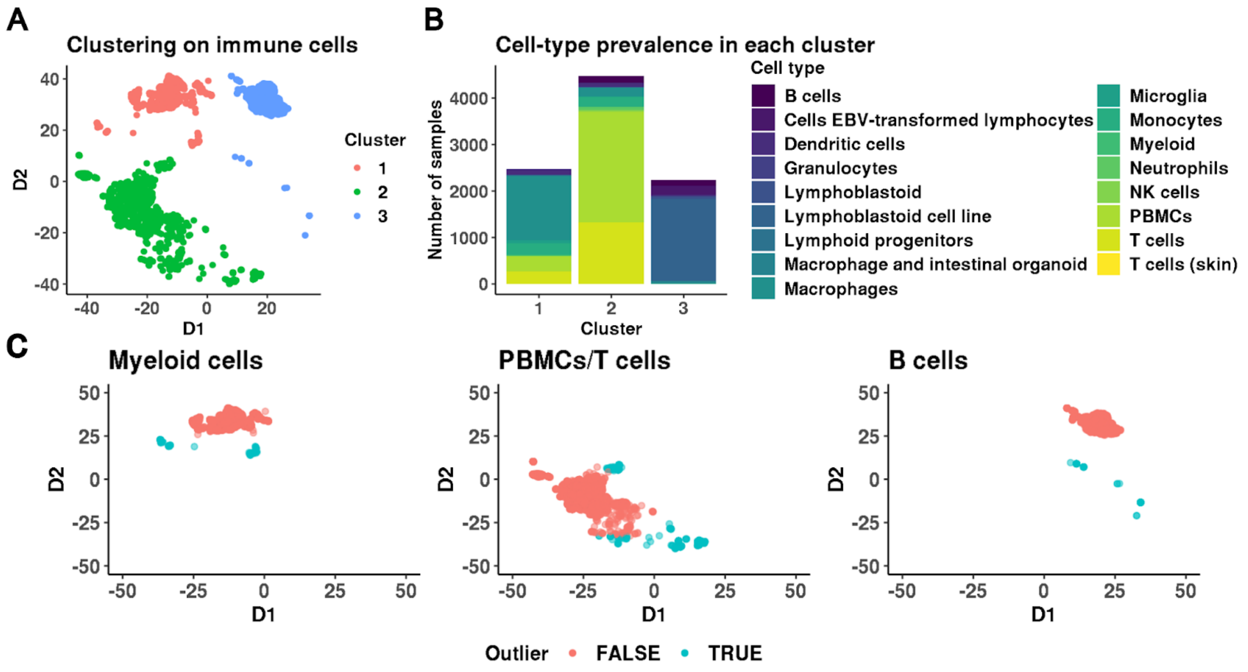


**Supp. Fig 2**

**(A)** We clustered the t-SNE projections of the xCell deconvolution scores of immune system samples using Gaussian mixture models to obtain three clusters with 2,472, 4,481 and 2,240 samples in clusters 1, 2 and 3 respectively. **(B)** Distribution of known tissue labels within each cluster; we classified each cluster based on the predominant cell type. **(C)** Identification of outliers in each immune context: We retained 88.87% of the samples in Cluster 1, 90.1% of the samples in Cluster 2 and 97.23% of the samples in Cluster 3.

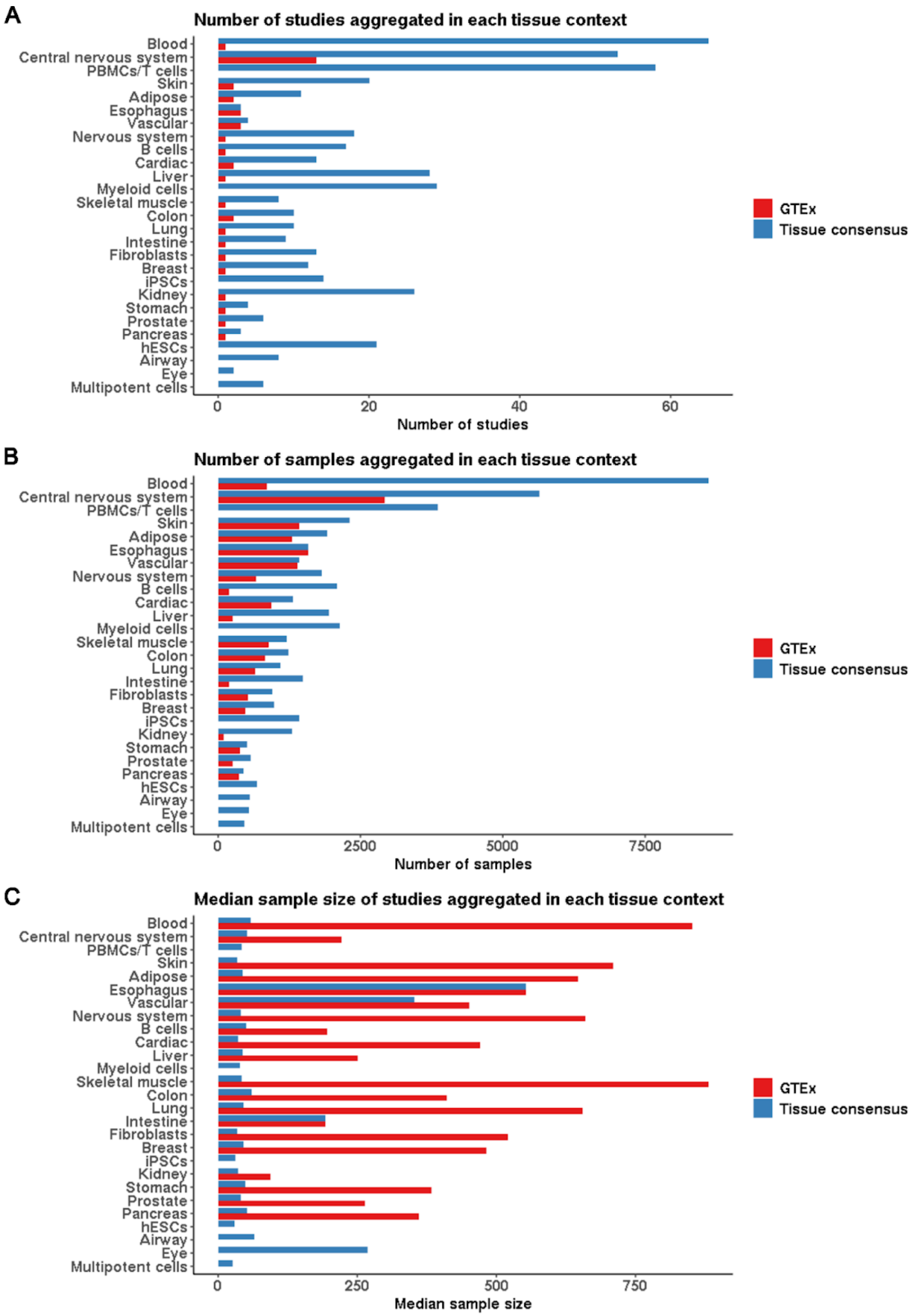
A
Number of studies aggregated in each tissue context
Blood
Central nervous system
PBMCs/T cells
Skin
Adipose
Esophagus
Vascular
Nervous system
B cells
Cardiac
Liver
Myeloid cells
Skeletal muscle
Colon
Lung
Intestine
Fibroblasts
Breast
iPSCs
Kidney
Stomach
Prostate
Pancreas
hESCs
Airway
Eye
Multipotent cells
0
20
40
60
Number of studies
GTEx
Tissue consensus
B
Number of samples aggregated in each tissue context
0
2500
5000
7500
Number of samples
C
Median sample size of studies aggregated in each tissue context
0
250
500
750
Median sample size

**Supp. Fig 3**

**(A)** Number of studies used in inferring tissue-specific GTEx and tissue-specific consensus networks across 27 contexts. **(B)** The number of samples from each tissue context used in the inference of tissue-specific GTEx and consensus networks. For 5 tissue contexts including hESCs, iPSCs, Eye, Airway, and Multipotent cells no samples were found in GTEx. **(C)** The median sample size of each study was aggregated to infer tissue-specific GTEx and consensus networks.

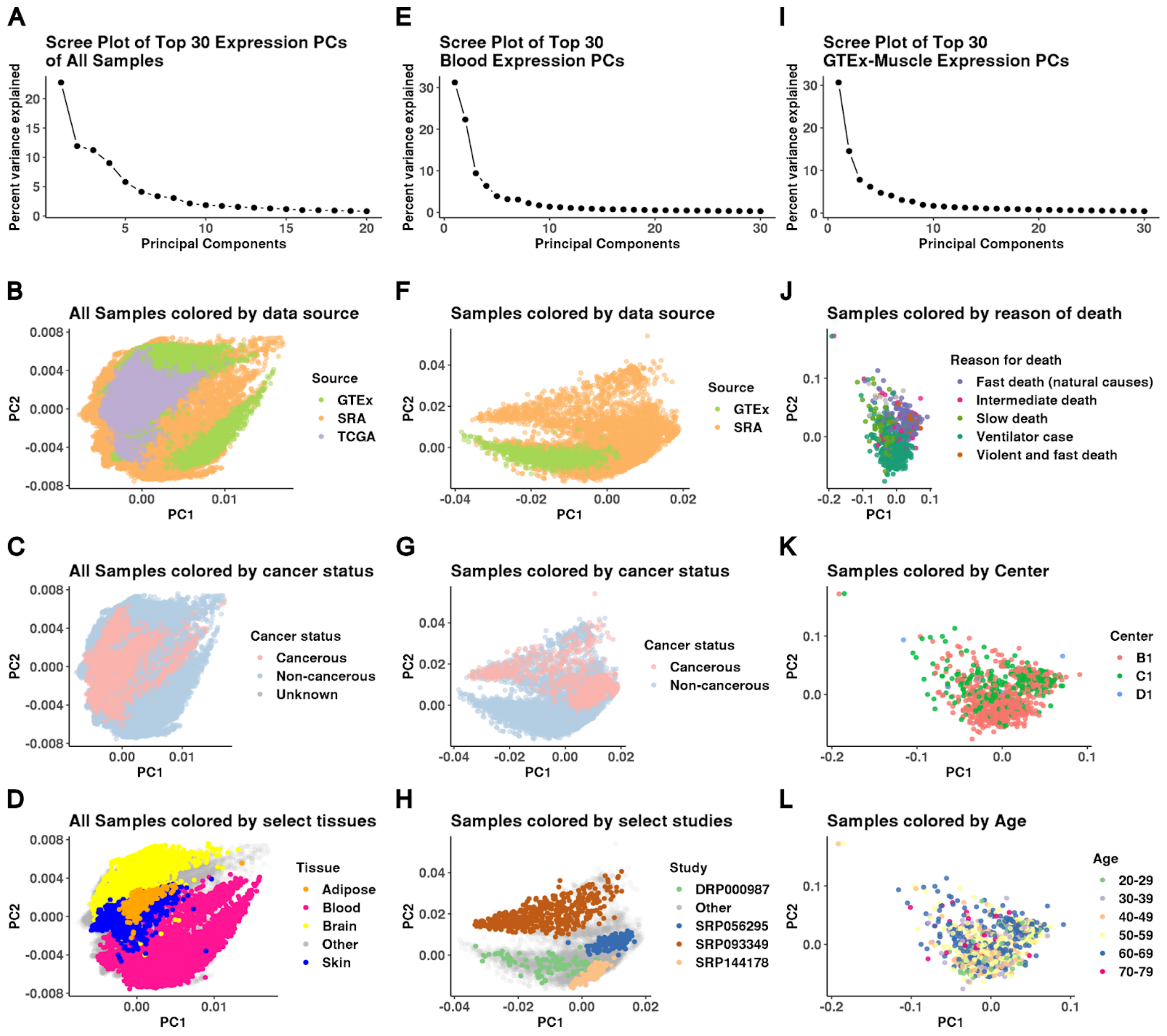


**Supp. Fig 4**

**(A)** Scree plot of top 30 PCs computed on gene expression from 95,280 samples. The top 10 PCs explain 75.2% of the total variance in gene expression. **(B)** Samples cluster by data source when projected onto the 2D space described by the top 2 PCs. **(C)** Cancerous samples cluster together when projected onto the 2D space described by the top 2 PCs. **(D)** Samples cluster by tissue label when projected onto the 2D space described by the top 2 PCs. **(E)** Scree plot of top 30 PCs computed on gene expression from 10,363 samples belonging to the blood tissue context. The top 7 PCs account for 80% of the variance in gene expression. **(F)** Blood samples from GTEx cluster together when expression data is projected onto the 2D PCA space. **(G)** PC2 captures the variance in gene expression driven by cancer status. **(H)** Samples from studies DRP00987, SRP056295, SRP093349, and SRP144178 cluster together when projected onto 2D PCA space. **(I)** Scree plot of 30 PCs computed on gene expression of GTEx skeletal muscle samples. The top 10 PCs account for 77.35% of the variance in gene expression. **(J)** Skeletal muscle samples cluster by reason of death when projected onto the 2D space described by the top 2 PCs. **(K)** Projecting skeletal muscle samples onto 2D PCA space results in weak clustering of samples by the processing center. **(L)** Projecting skeletal muscle samples onto 2D PCA space does not result in a strong clustering of samples by age.

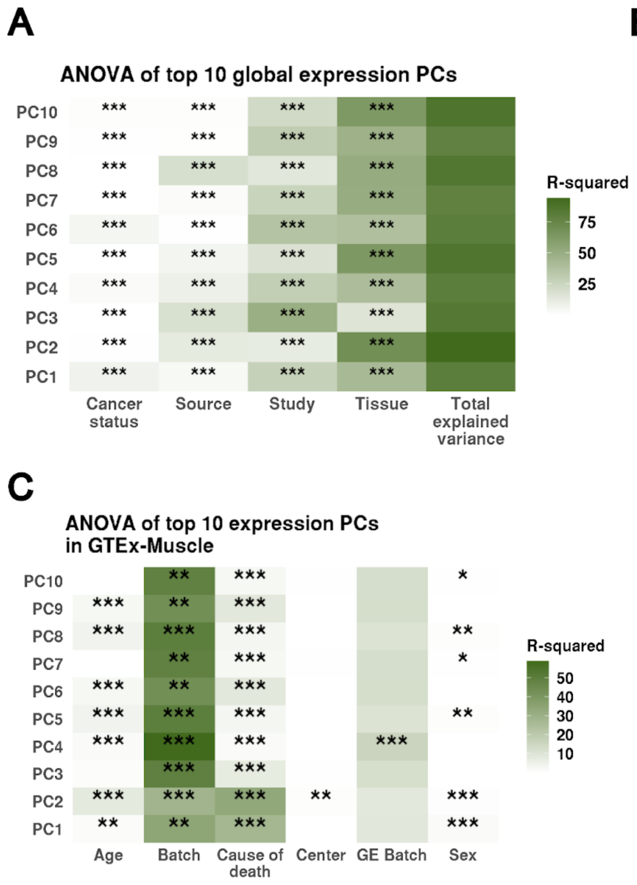


**Supp. Fig 5**

R-squared (Percent of variance explained) of PCs computed across samples by sample characteristics, as well as the significance of the p-value corresponding to F-statistic for each explanatory variable obtained by ANOVA. *** : $p < 1e-3$, ** : $1e-3 < p \leq 1e-2$, *: $1e-2 < p < 5e-2$. Calculated across **(A)** all samples, **(B)** all blood samples, and **(C)** GTEx skeletal muscle samples.

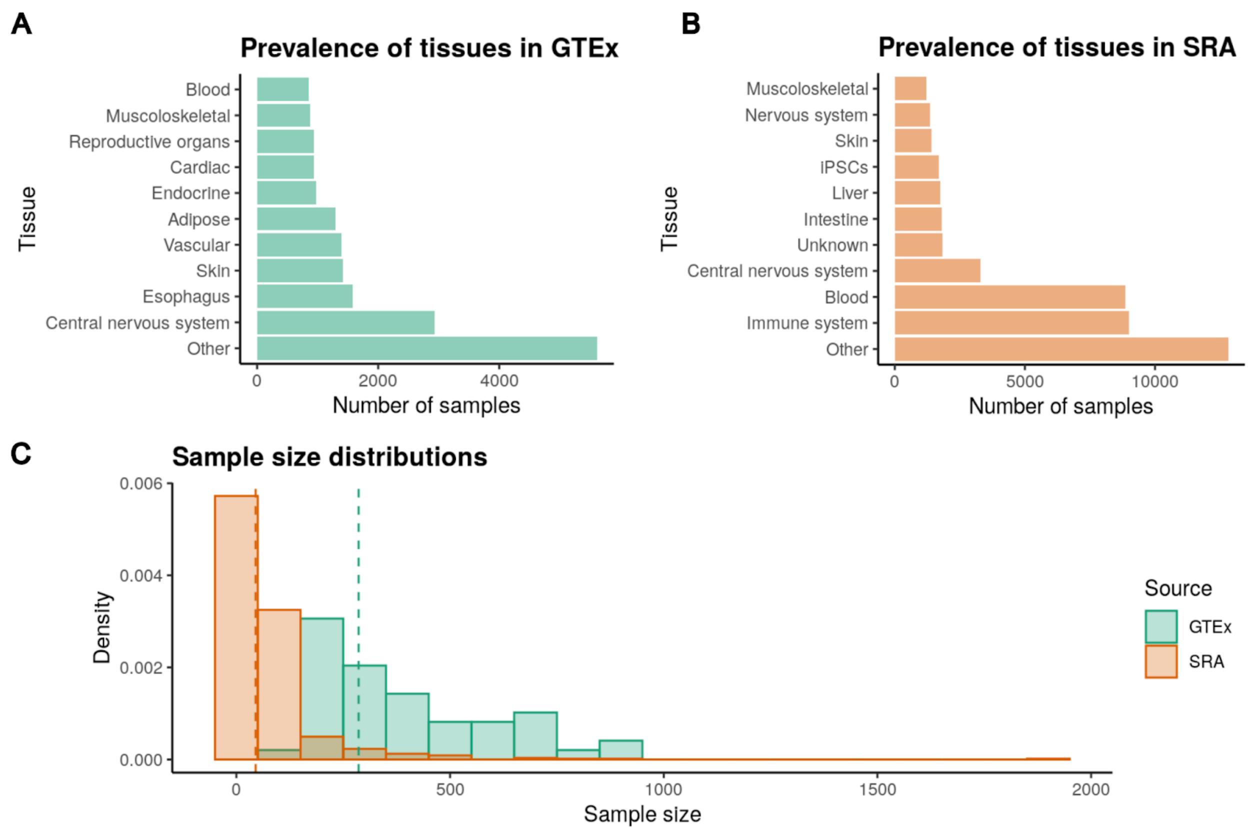


**Supp. Fig 6**

**(A)** Tissue composition of GTEx data split for top 10 tissues by sample size and the number of samples which do not belong to one of the top 10 tissues. **(B)** Tissue composition of SRA data split for the top 10 tissues by sample size and the number of samples which do not belong to one of the top 10 tissues. 1,843 samples do not have a known tissue type. **(C)** Sample are grouped by tissue in GTEx and study in SRA. The median sample size of GTEx tissues is 286 and the median sample of SRA studies is 45.

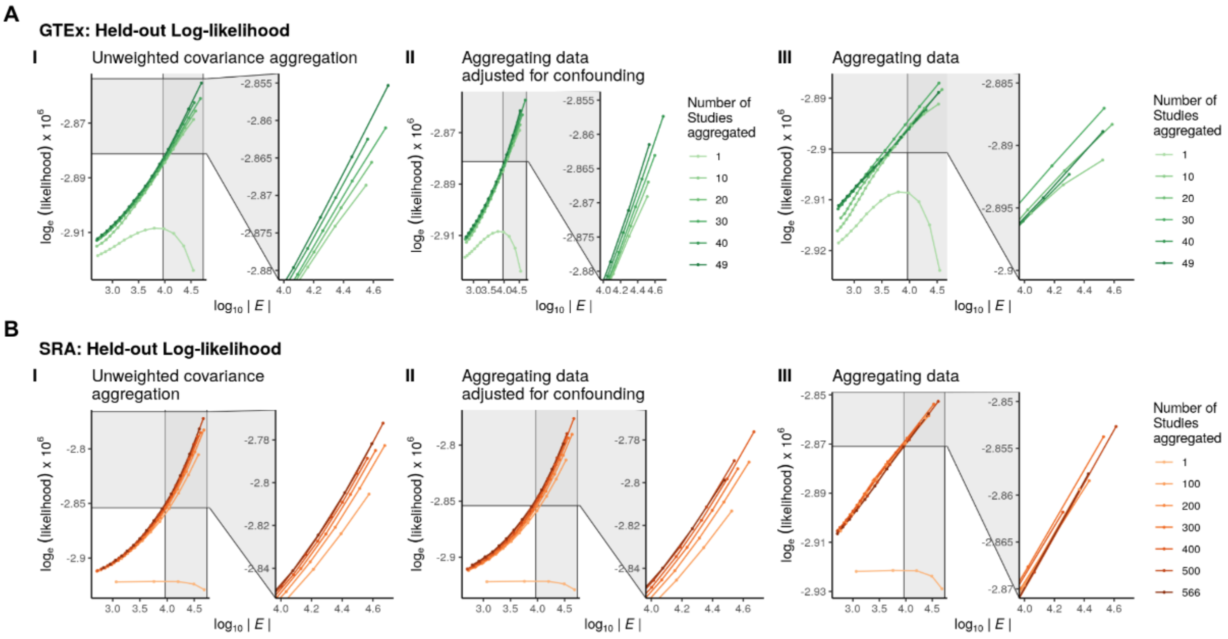


**Supp. Fig 7**

**(A)** Impact of the number of GTEx tissues aggregated on the held-out log-likelihood estimated using SRA data for networks with densities between $10^3$ and $10^5$ when data is aggregated using three data aggregation strategies. **(B)** Impact of the number of SRA studies aggregated on the held-out log-likelihood estimated using GTEx data for networks with densities between $10^3$ and $10^5$ when data is aggregated using three data aggregation strategies.

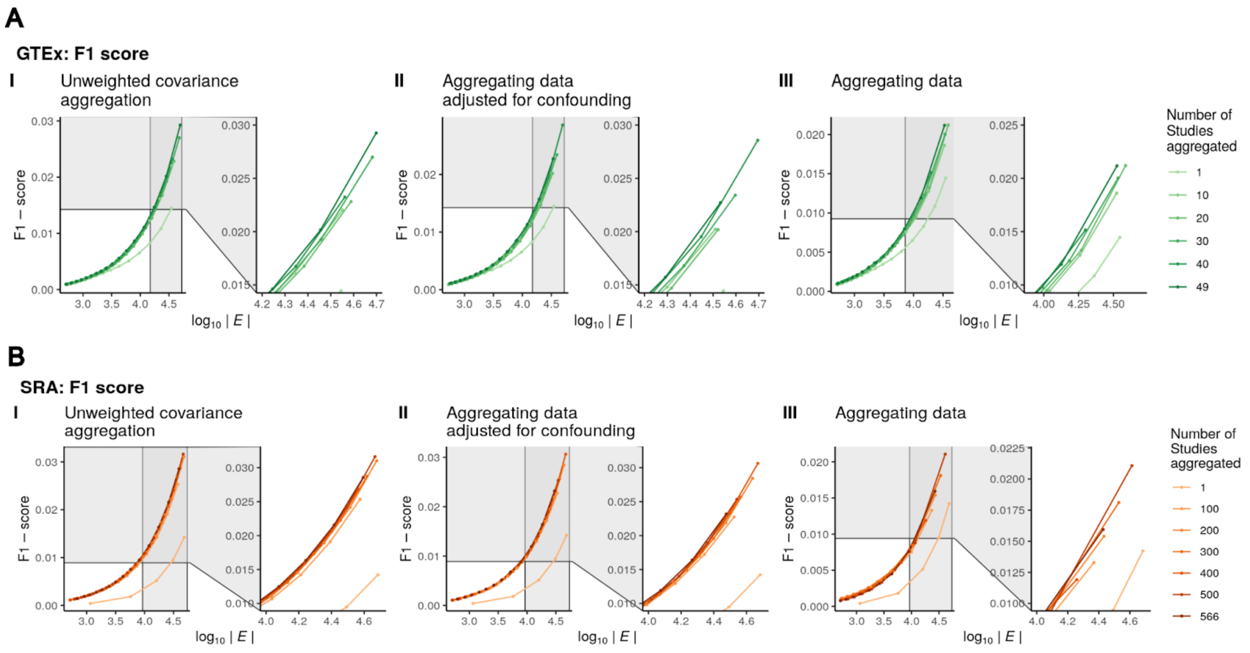


**Supp. Fig 8**

**(A)** Impact of data aggregation on F1-scores of GTEx networks compared to consensus pathways across data aggregation strategies. **(B)** Impact of data aggregation on F1-scores of SRA networks compared to consensus pathways across data aggregation strategies.

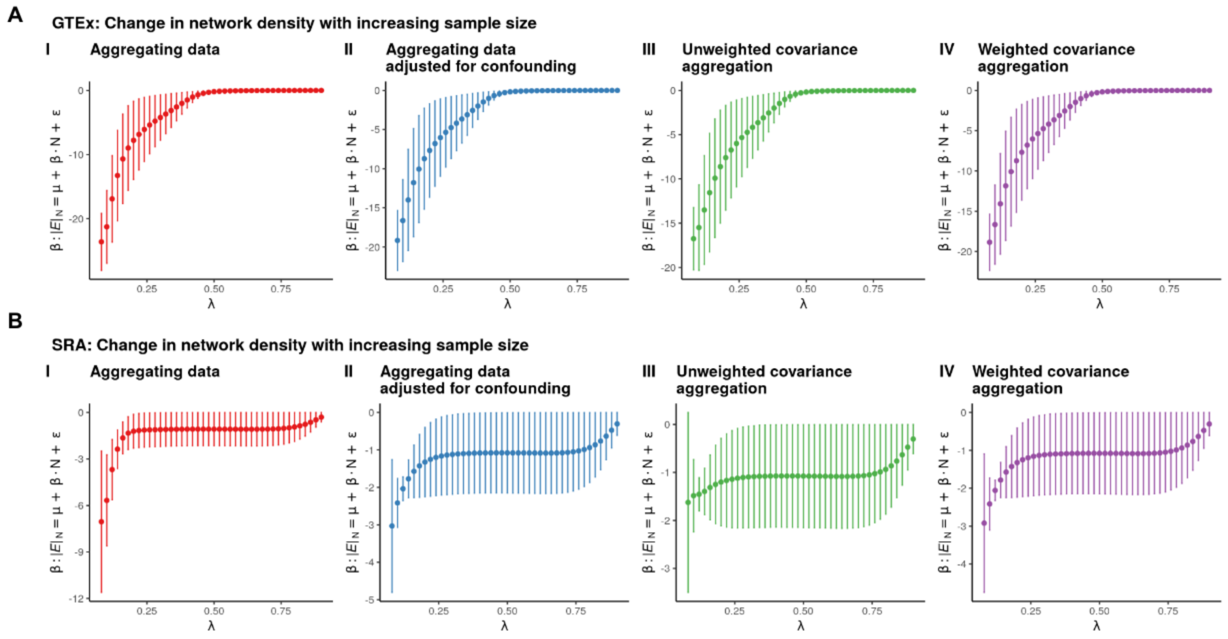


**Supp. Fig 9**

Regression estimate (β) and standard errors of the effect of sample size (N) on network density across varying penalization parameters $\lambda$ for (**A**) GTEx networks and (**B**) SRA networks inferred from **(I)** Aggregating normalized data prior to PC-based data correction **(II)** Aggregating PC-corrected data **(III)** Unweighted aggregation of covariance matrices **(IV)** Weighted aggregation covariance matrices.

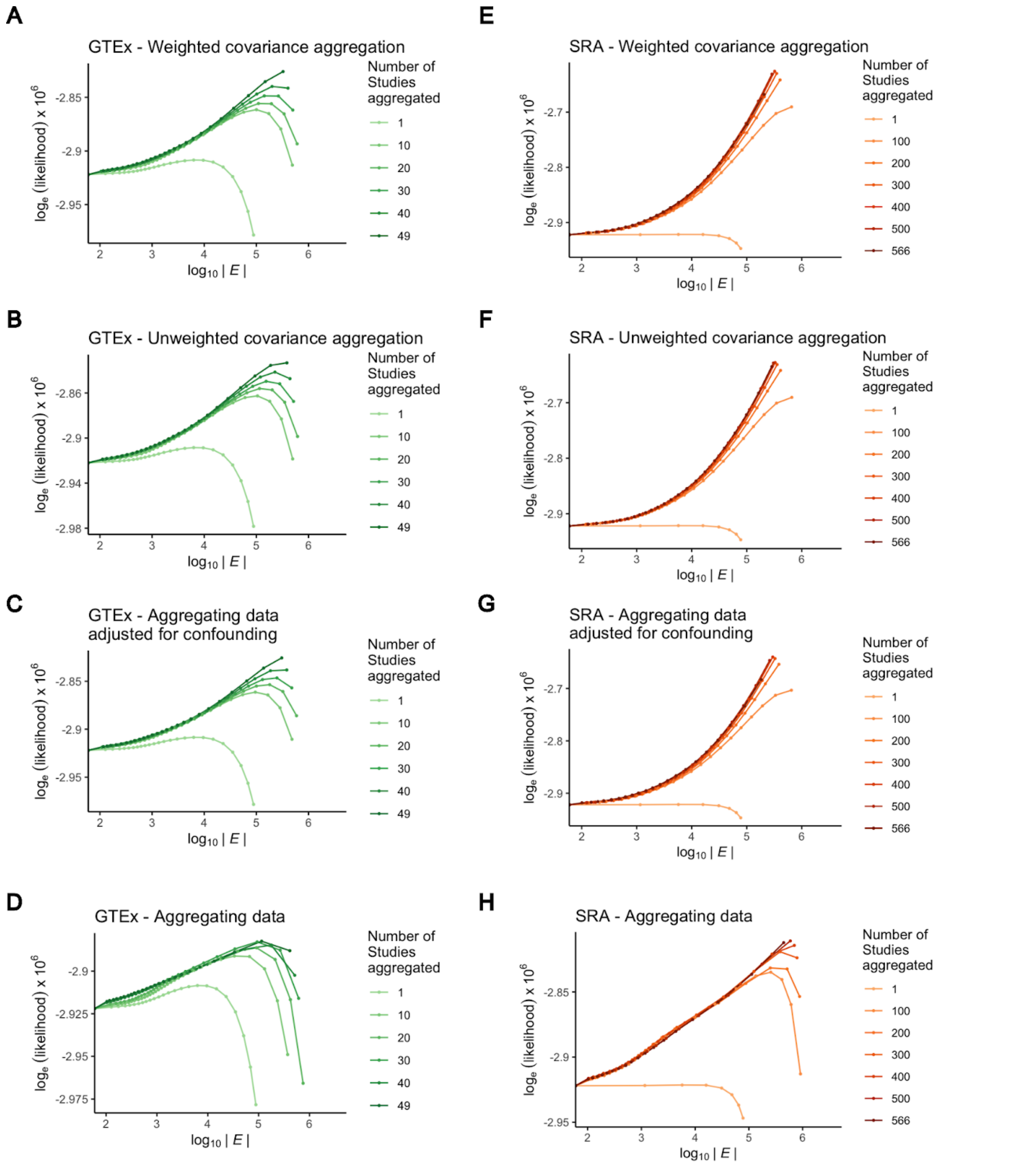


**Supp. Fig 10**

Held-out log-likelihood of networks inferred by sequential data aggregation of **(A - D)** GTEx studies 10 at a time and **(E-H)** SRA studies 100 at a time using different aggregation strategies increased with density for networks with fewer than 1e6 edges. However, denser networks (>

1e6 edges) which have a greater log-likelihood for the training data resulted in a lower held-out log-likelihood implying overfitting.

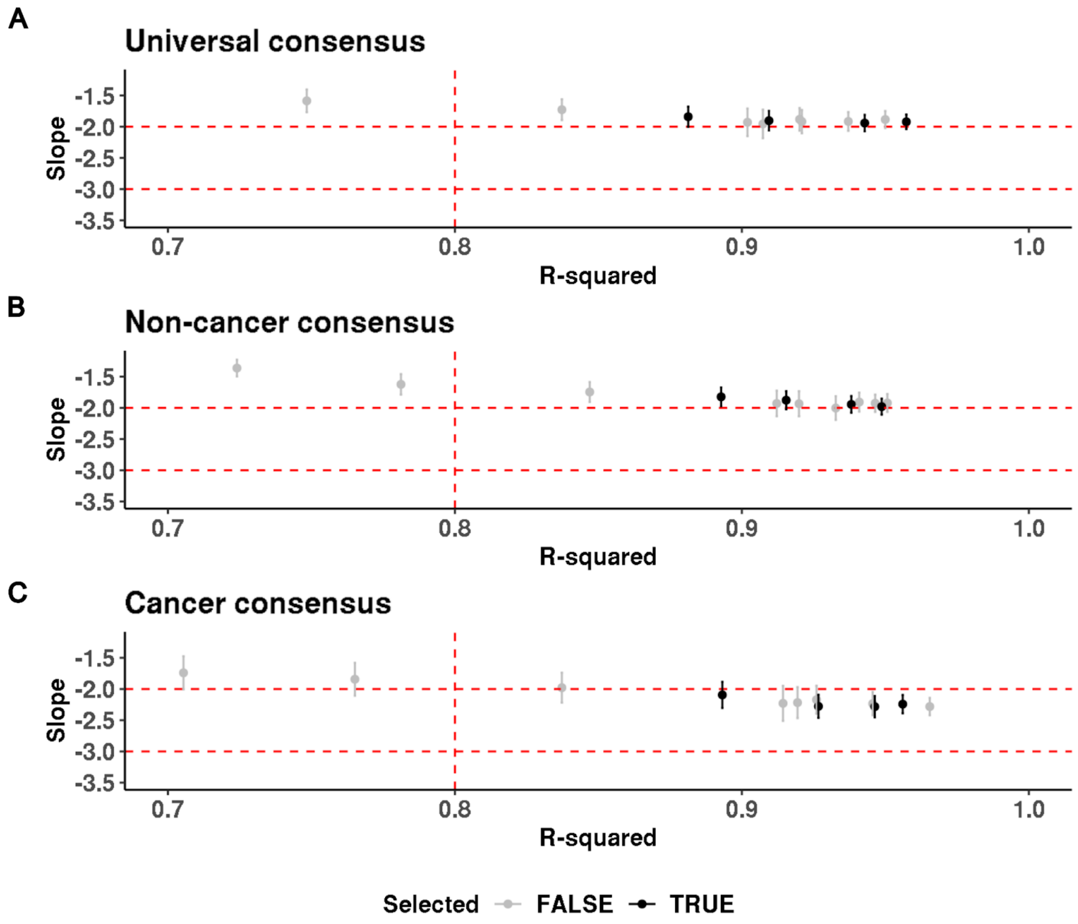


**Supp. Fig 11**

Estimate of slope vs $R^2$ obtained from fitting a linear model between the log empirical degree distribution and log degree. (A) Universal consensus networks, penalization parameters $\lambda$ = 0.14, 0.16, 0.18, and 0.20 were selected. **(B)** Non-cancer consensus networks, penalization parameters $\lambda$ = 0.14, 0.16, 0.18, and 0.20 were selected. **(C)** Cancer consensus networks, penalization parameters $\lambda$ = 0.20, 0.22, 0.24, and 0.26 were selected.

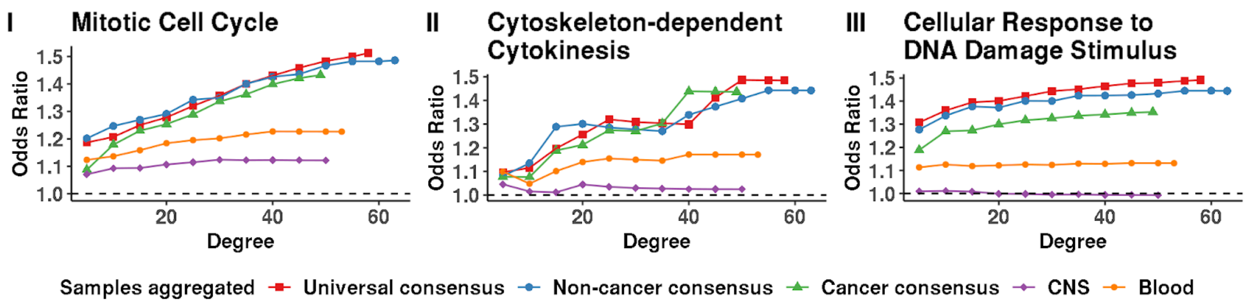


**Supp. Fig 12**

**(A)** The odds ratio of finding genes involved in ubiquitous biological processes including **(I)** mitotic cell cycle (GO:0000278), **(II)** cytoskeleton-dependent cytokinesis (GO:0061640), and **(III)** cellular response to DNA (GO:0006974) in groups of network nodes from three consensus networks, universal (λ = 0.18, 7087 edges), non-cancer (λ = 0.18, 7355 edges), and cancer (λ = 0.24, 7552 edges) compared to context-specific networks inferred corresponding to blood (λ = 0.24, 7283 edges), and CNS (λ = 0.28, 8430 edges) selected by progressively increasing degree (X-axis).

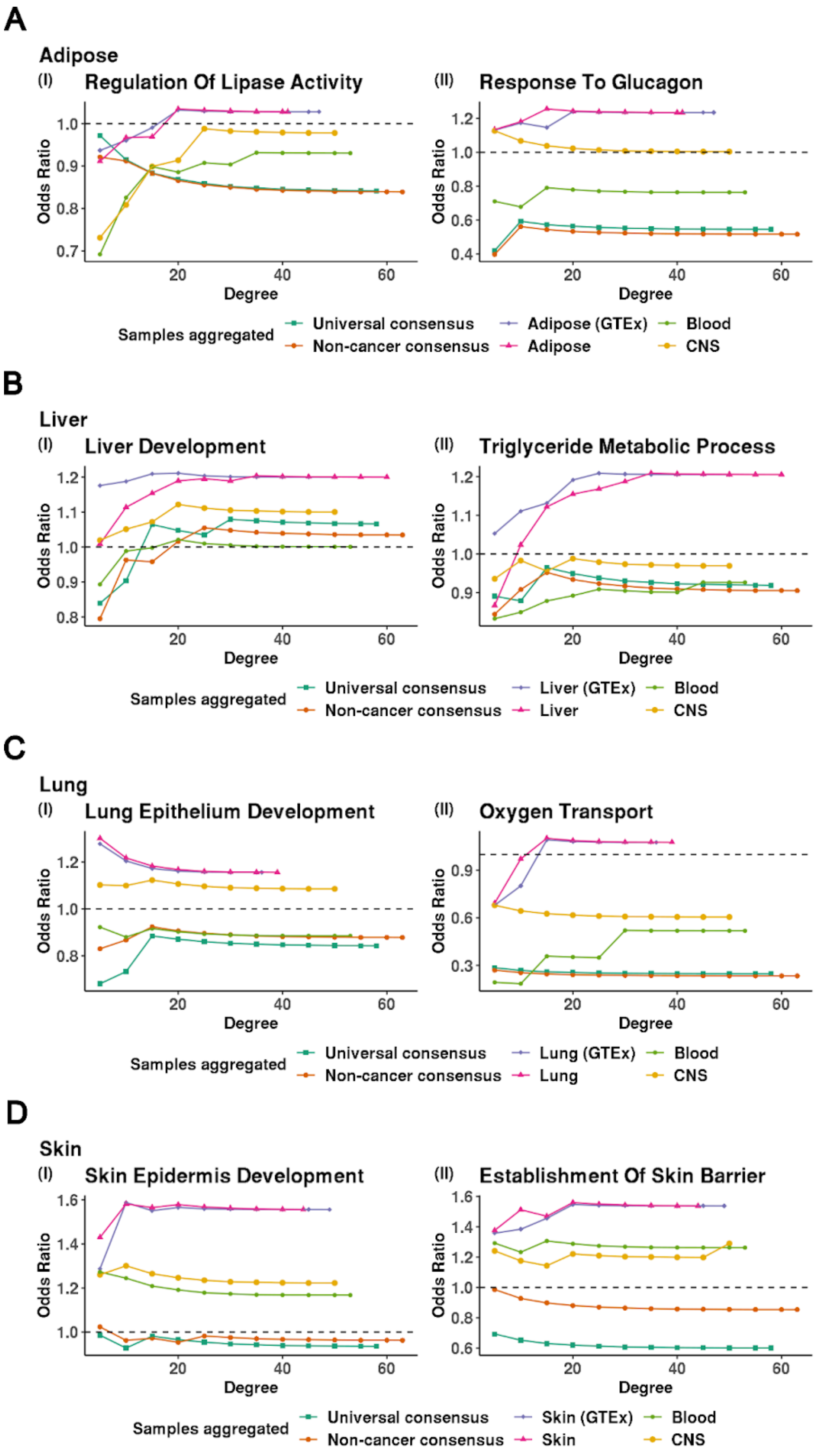
A
Adipose
(I) Regulation Of Lipase Activity
(II) Response To Glucagon
Odds Ratio
Degree
Samples aggregated
Universal consensus
Non-cancer consensus
Adipose (GTEx)
Adipose
Blood
CNS
B
Liver
(I) Liver Development
(II) Triglyceride Metabolic Process
Odds Ratio
Degree
Samples aggregated
Universal consensus
Non-cancer consensus
Liver (GTEx)
Liver
Blood
CNS
C
Lung
(I) Lung Epithelium Development
(II) Oxygen Transport
Odds Ratio
Degree
Samples aggregated
Universal consensus
Non-cancer consensus
Lung (GTEx)
Lung
Blood
CNS
D
Skin
(I) Skin Epidermis Development
(II) Establishment Of Skin Barrier
Odds Ratio
Degree
Samples aggregated
Universal consensus
Non-cancer consensus
Skin (GTEx)
Skin
Blood
CNS

**Supp. Fig 13**

Enrichment of genes involved in GO processes that are tissue-specific among network genes selected with increasing thresholds of degree connectivity in concordant tissues **(A)** Adipose ($\lambda$ = 0.26, 7573), Adipose (GTEx) ($\lambda$ = 0.28, 7192), **(B)** Liver ($\lambda$ = 0.24, 7573), Liver (GTEx) ($\lambda$ = 0.38, 8027), **(C)** Lung ($\lambda$ = 0.28, 8313), Lung (GTEx) ($\lambda$ = 0.30, 7304), **(D)** Skin ($\lambda$ = 0.26, 7567), Skin (GTEx) ($\lambda$ = 0.28, 6428) compared to two consensus networks consensus networks, universal ($\lambda$ = 0.18, 7087 edges), non-cancer ($\lambda$ = 0.18, 7355 edges), as well as two discordant context-specific networks corresponding to blood ($\lambda$ = 0.24, 7283 edges), and CNS ($\lambda$ = 0.28, 8430 edges)selected by progressively increasing degree (X-axis).

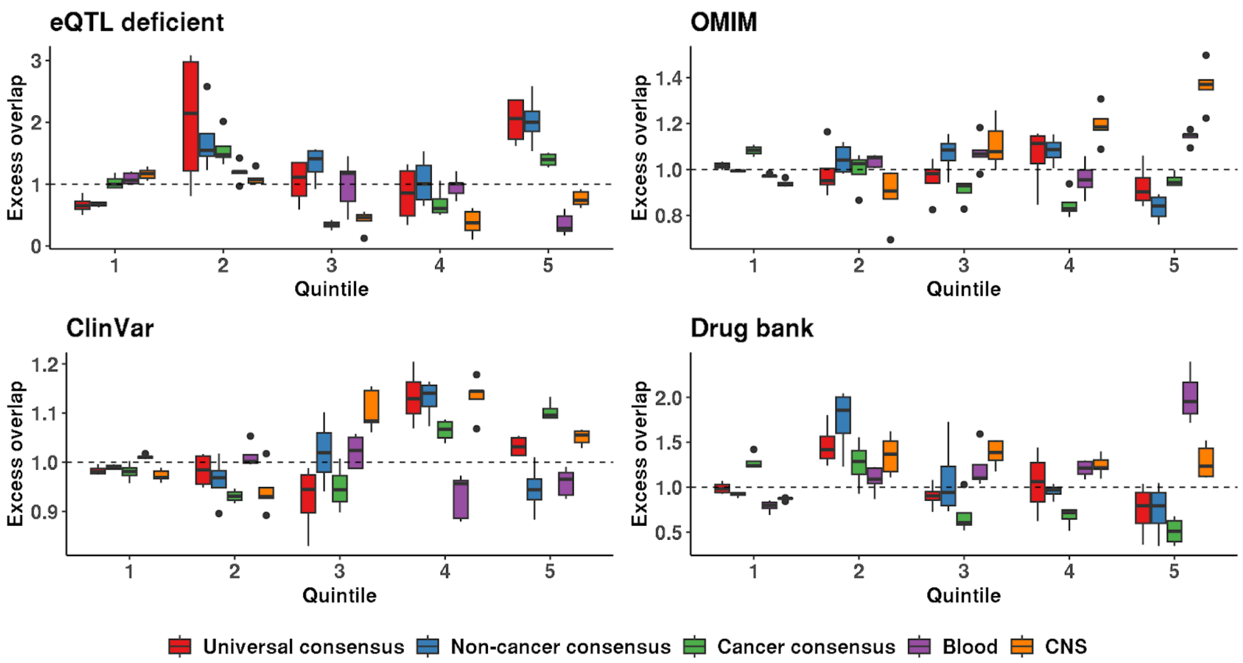


**Supp. Fig 14**

Distribution of the excess overlap of functional gene sets for network nodes binned by the number of neighbors (degree) corresponding to universal consensus networks (λ = 0.14, 0.16, 0.18, 0.20), non-cancer consensus network (λ = 0.14, 0.16, 0.18, 0.20), cancer consensus networks (λ = 0.20, 0.22, 0.24, 0.26), blood network (λ = 0.18, 0.20, 0.22, 0.24, 0.26), and CNS network (λ = 0.24, 0.26, 0.28, 0.30, 0.32). Nodes with no neighbors are grouped (Quintile 1). Nodes with non-zero neighbors are split based on the quartile they belong to (Quintiles 2-5). **(A)** The excess overlap of 604 eQTL deficient genes with no significant variant-gene association in all 48 tissues in GTEx v7 single-tissue cis-eQTL data. **(B)** 2,266 genes deposited in the Online Mendelian Inheritance in Man (OMIM). **(C)** 5,428 ClinVar genes with a pathogenic or likely pathogenic variant with no conflict among studies. **(D)** 373 Drug bank genes whose protein products are human targets of FDA-approved drugs with known mechanisms of action.

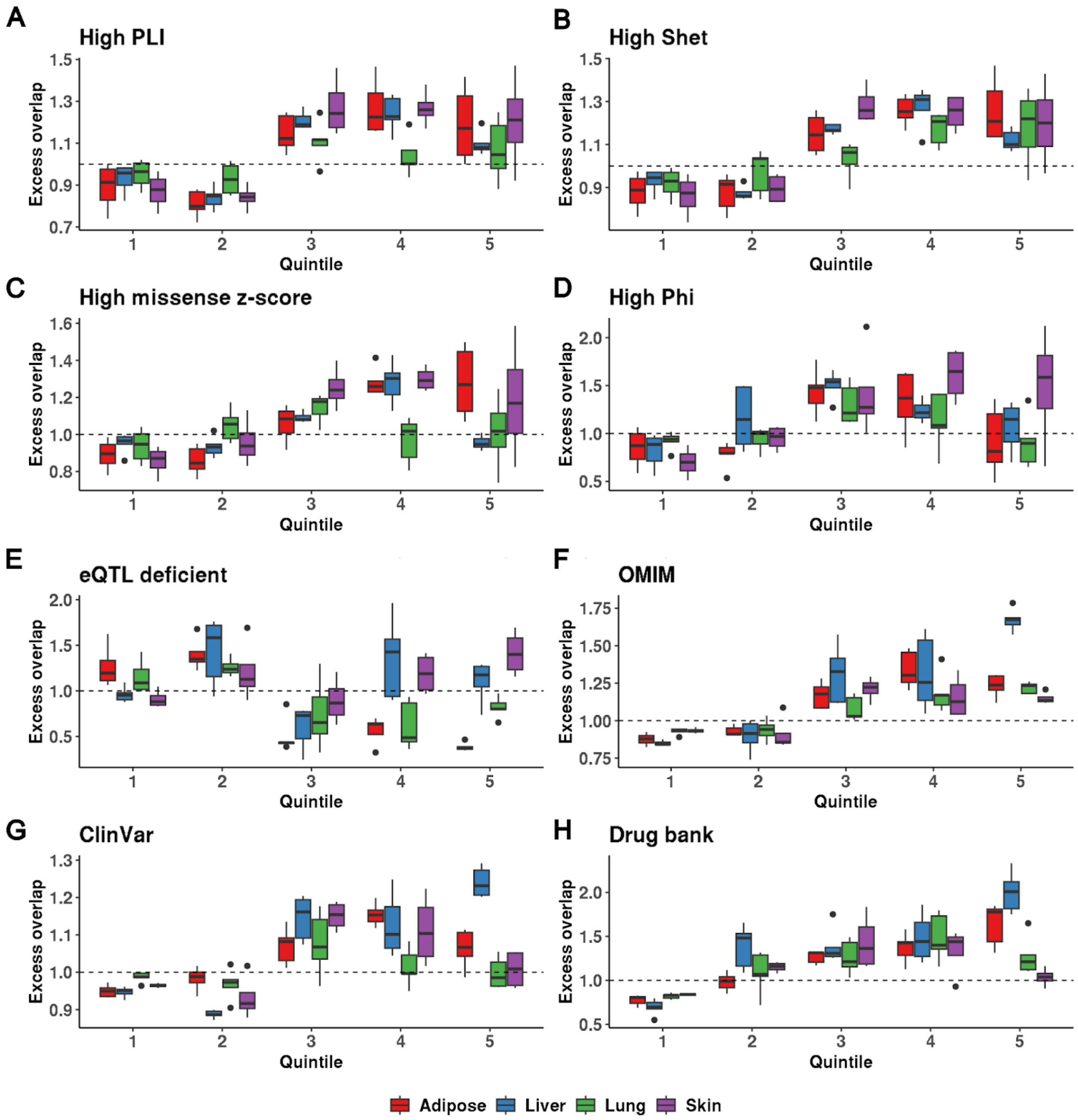


**Supp. Fig 15**

Distribution of the excess overlap of evolutionarily conserved and functional gene sets for network nodes binned by the number of neighbors (degree) derived from Adipose (λ = 0.20 - 0.28), Liver (λ = 0.20 - 0.28), Lung (λ = 0.24 - 0.32), and Skin (λ = 0.22 - 0.28) networks. Nodes with no neighbors are grouped (Quintile 1). Nodes with non-zero neighbors are split based on the quartile they belong to (Quintiles 2-5). The excess overlap of **(A)** 3,104 loss-of-function (LoF) genes with pLI > 0.9, i.e., strongly depleted for protein-truncating variants **(B)** 2,853 genes with a $S_{het} > 0.1$ **(C)** 1,440 genes strongly depleted for missense mutations (high missense z-score) **(D)** 588 genes with a Phi-score > 0.95 **(E)** 604 genes with no significant cis-eQTL in GTEx **(F)** 2,266 disease genes deposited in OMIM **(G)** 5,428 ClinVar genes with a pathogenic or likely pathogenic variant **(H)** 373 DrugBank genes whose protein products are human targets of FDA-approved drugs with known mechanisms of action.

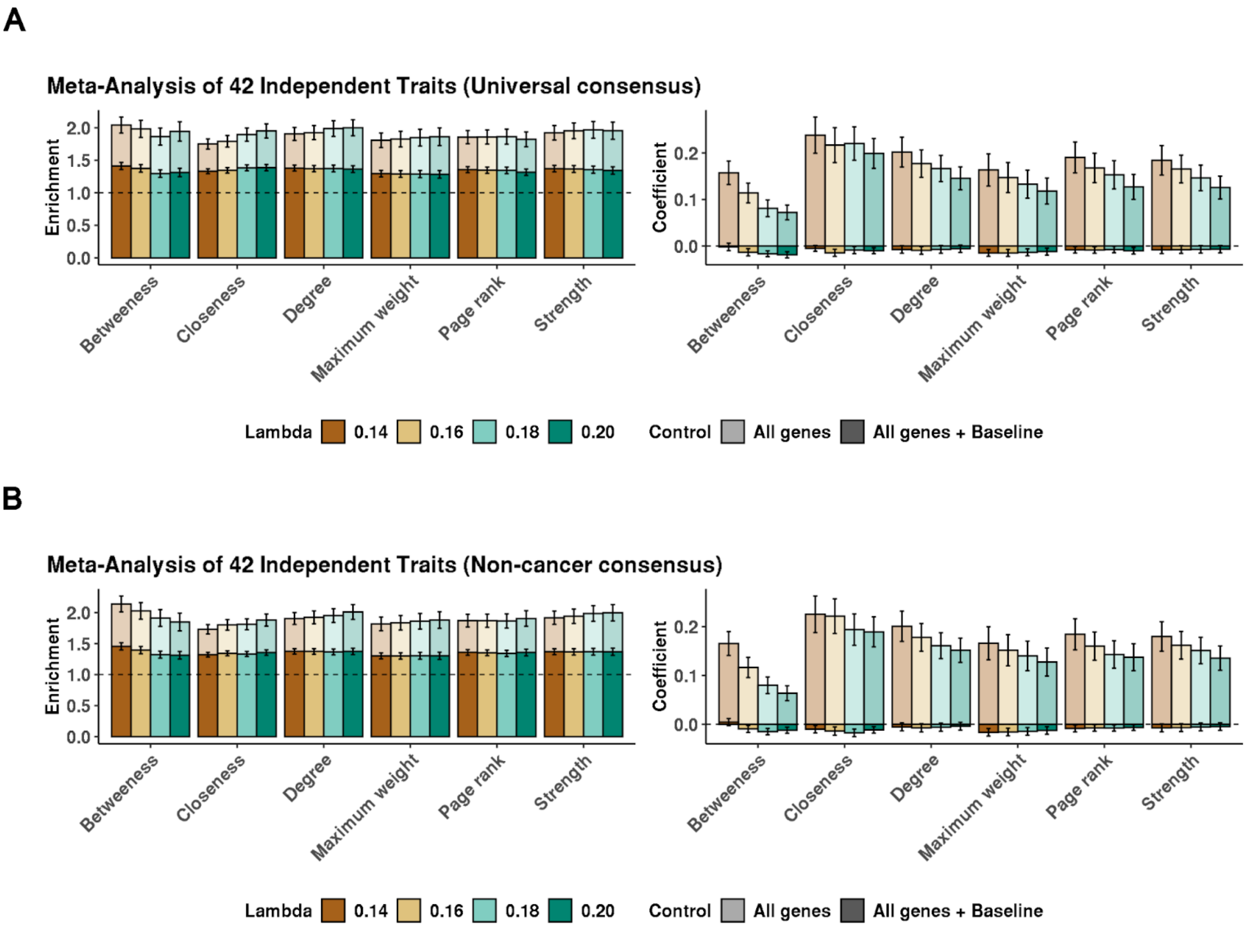


**Supp. Fig 16**

Estimates of the heritability enrichment and the coefficient $\tau^*$, meta-analyzed across 42 independent traits for annotations corresponding to 6 measures of network centrality derived from **(A)** universal consensus networks for a range of L1-penalty parameter between 0.14 - 0.20 and network density between 17,320 - 4,852 and **(B)** non-cancer consensus networks for a range of L1-penalty parameter between 0.14-0.20 and network density between 17,476 - 5,144.

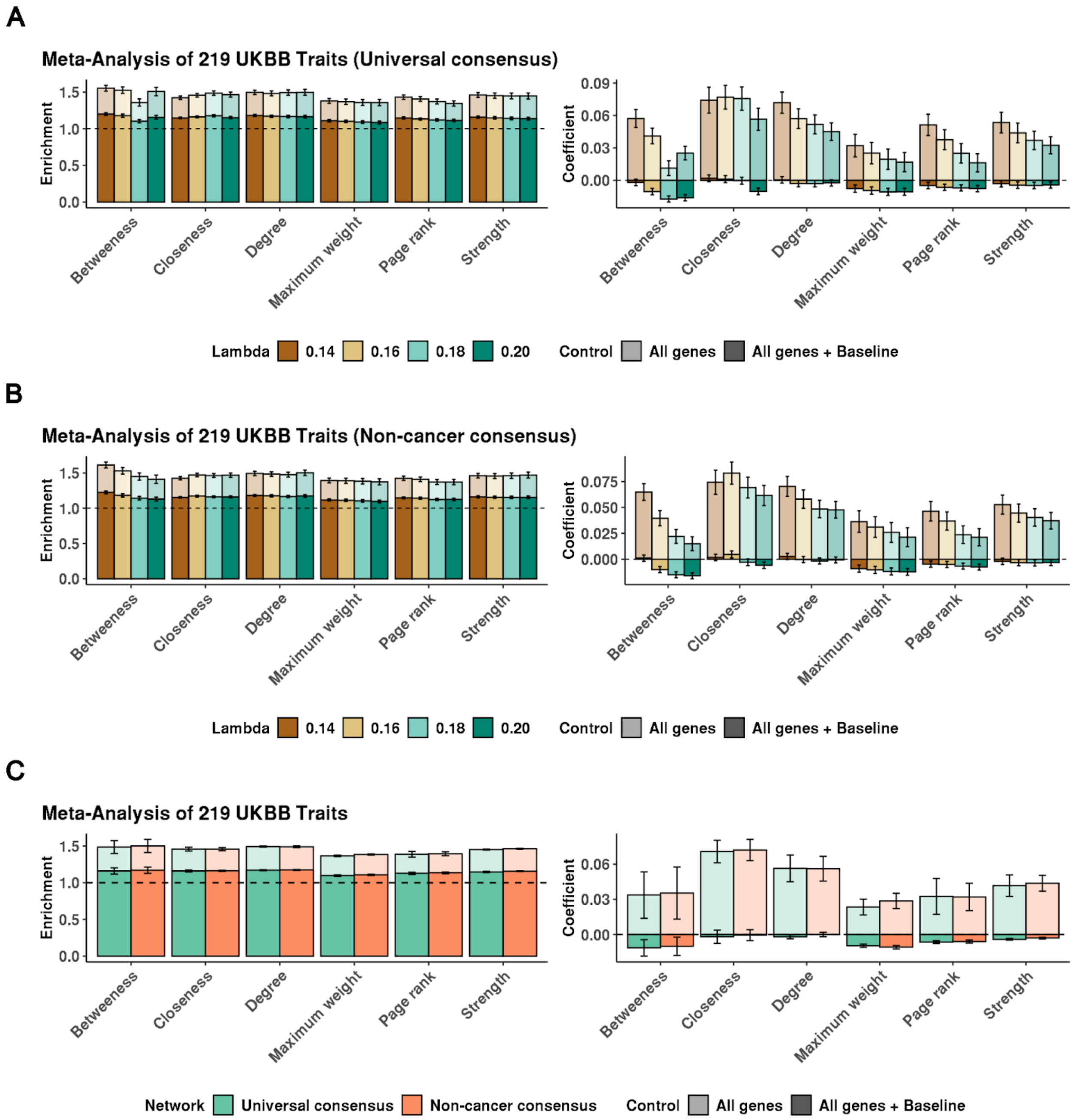


**Supp. Fig 17**

Estimates of the heritability enrichment and the coefficient $\tau^*$, meta-analyzed across 219 UKBB traits for annotations corresponding to 6 measures of network centrality derived from (A) universal consensus networks for a range of L1-penalty parameters between 0.14 - 0.20 and network density between 17,320 - 4,852. **(B)** from non-cancer consensus networks for a range of

L1-penalty parameters between 0.14 - 0.20 and network density between 17,476 - 5,144. **(C)** Comparison of the average heritability enrichment and coefficient values between networks when conditioning on the all-genes or baseline-LD annotations.

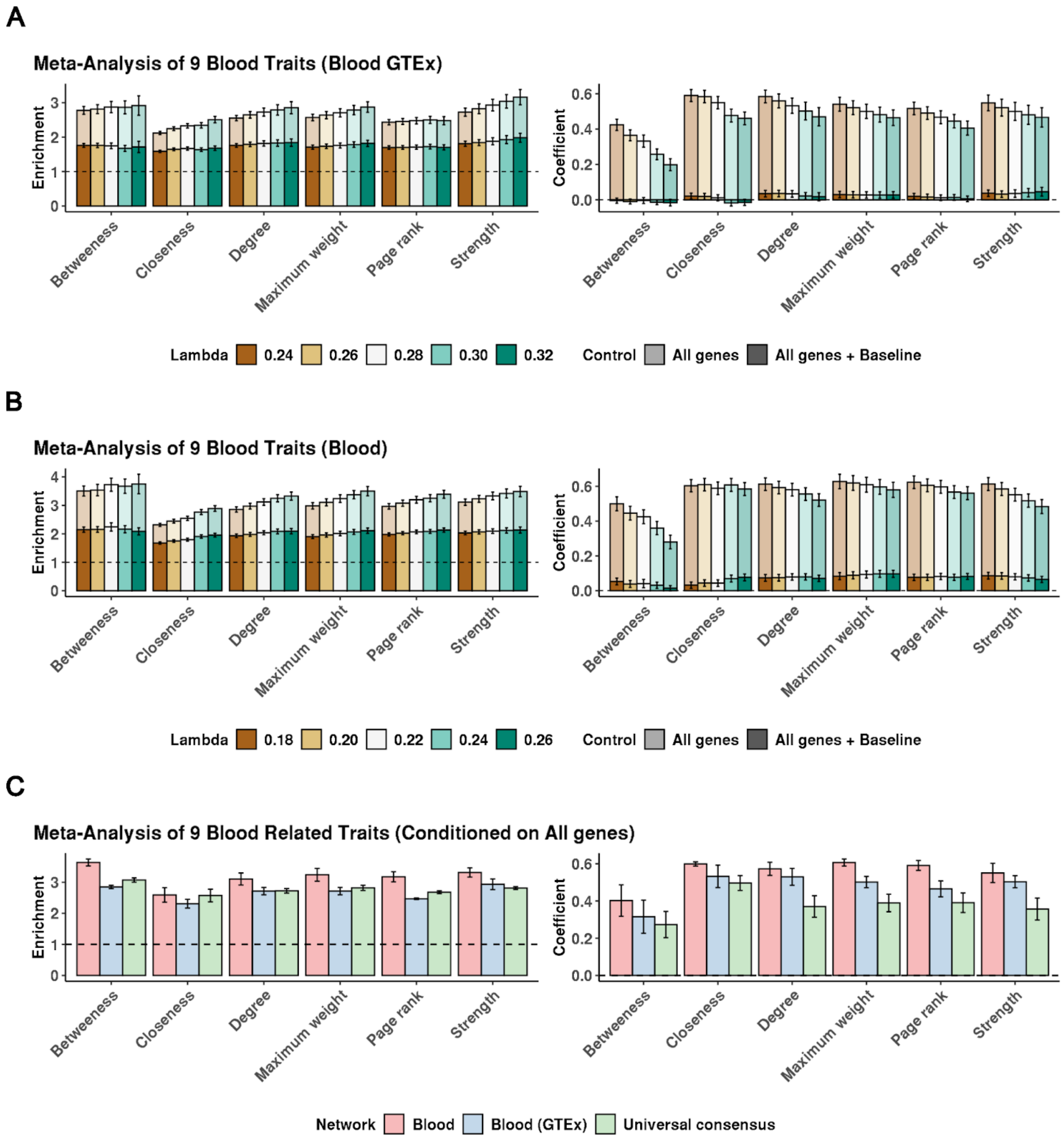


**Supp. Fig 18**

Estimates of the heritability enrichment and the coefficient, $\tau^*$, meta-analyzed across 9 blood traits for annotations corresponding to 6 measures of network centrality derived from **(A)** blood GTEx networks for a range of L1-penalty parameter between 0.24 - 0.32 and network density between 17,320 - 4,852 or **(B)** blood consensus networks for a range of L1-penalty parameters between 0.18 - 0.26 and network density between 18,153 - 5,636. **(C)** Comparison of the heritability enrichment and coefficient averages between the blood consensus networks ($\lambda$ = 0.18 - 0.26) blood GTEx networks ($\lambda$ = 0.24 - 0.32), and universal consensus networks ($\lambda$ = 0.14 - 0.20).

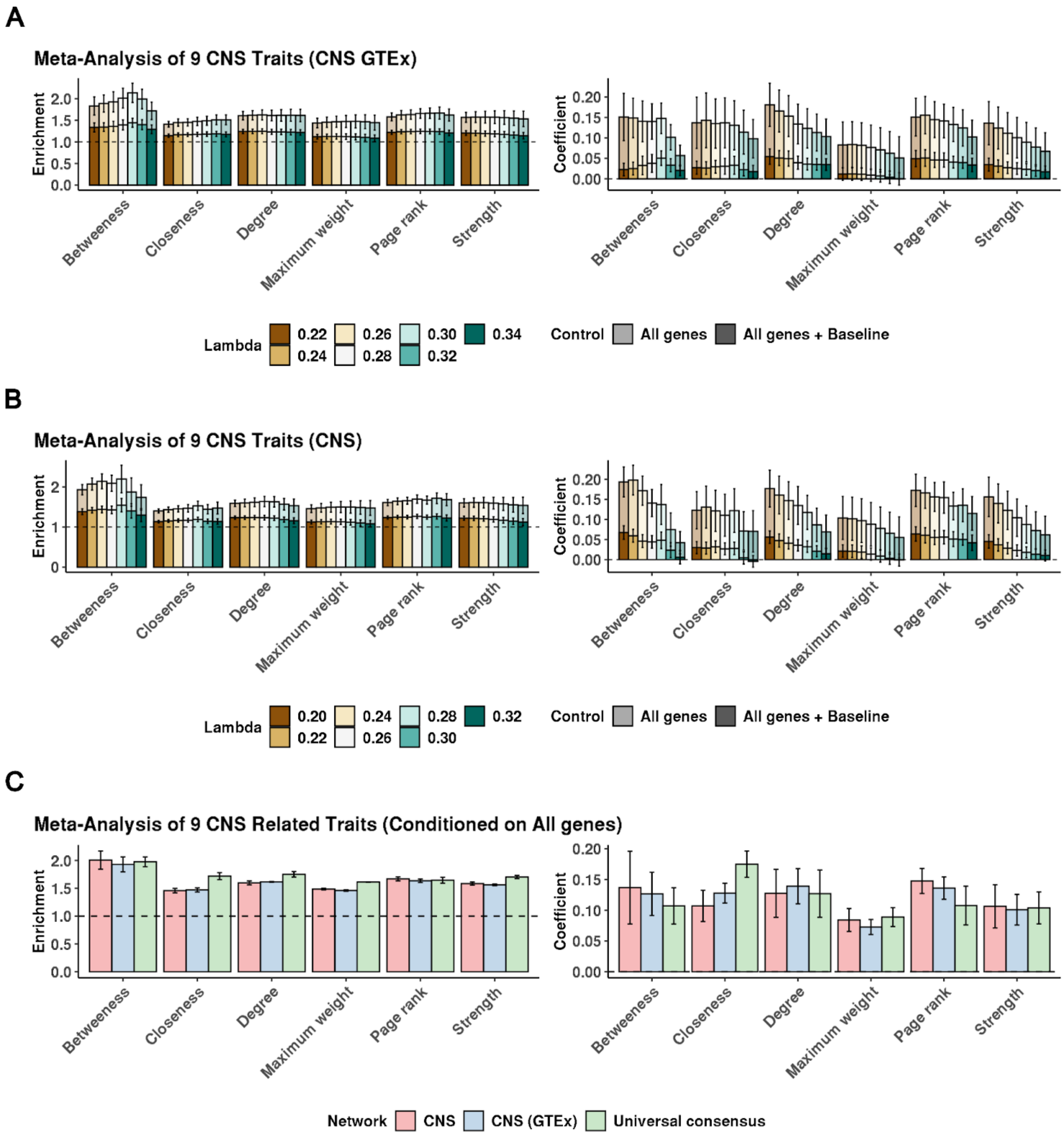


**Supp. Fig 19**

Estimates of the heritability enrichment and the coefficient, $\tau^*$, meta-analyzed across 9 CNS traits for annotations corresponding to 6 measures of network centrality derived from **(A)** CNS GTEx networks for a range of L1-penalty parameter between 0.22 - 0.34 and network density between 32,072 -7,807 or **(B)** CNS consensus networks for a range of L1-penalty parameters between 0.20 - 0.32 and network density between 29,710 - 4,783. **(C)** Comparison of heritability enrichment and coefficient across CNS consensus ($\lambda$ = 0.20 - 0.32), CNS GTEx ($\lambda$ = 0.22 - 0.34), and universal consensus networks ($\lambda$ = 0.14 - 0.20).
